# Cortical microstructural gradients capture memory network reorganization in temporal lobe epilepsy

**DOI:** 10.1101/2022.10.31.513891

**Authors:** Jessica Royer, Sara Larivière, Raul Rodriguez-Cruces, Donna Gift Cabalo, Shahin Tavakol, Hans Auer, Bo-yong Park, Casey Paquola, Jonathan Smallwood, Elizabeth Jefferies, Lorenzo Caciagli, Andrea Bernasconi, Neda Bernasconi, Birgit Frauscher, Boris C. Bernhardt

## Abstract

Temporal lobe epilepsy (TLE), one of the most common pharmaco-resistant epilepsies, is associated with pathology of paralimbic brain regions, particularly in the mesiotemporal lobe. Cognitive dysfunction in TLE is frequent, and particularly affects episodic memory. Crucially, these difficulties challenge the quality of life of patients, sometimes more than seizures, underscoring the need to assess neural processes of cognitive dysfunction in TLE to improve patient management. Our work harnessed a novel conceptual and analytical approach to assess spatial gradients of microstructural differentiation between cortical areas based on high-resolution MRI analysis. Gradients track region-to-region variations in intracortical lamination and myeloarchitecture, serving as a system-level measure of structural and functional reorganization. Comparing cortex-wide microstructural gradients between 21 patients and 35 healthy controls, we observed a contracted gradient in TLE driven by reduced microstructural differentiation between paralimbic cortices and the remaining cortex with marked abnormalities in ipsilateral temporopolar and dorsolateral prefrontal regions. Findings were replicated in an independent cohort. Using an independent *post mortem* dataset, we observed that *in vivo* findings reflected topographical variations in cortical lamination patterns, confirming that TLE-related changes in the microstructural gradient reflected increased proximity of regions with more dissimilar laminar structure. Disease-related transcriptomics could furthermore show specificity of our findings to TLE over other common epilepsy syndromes. Finally, microstructural dedifferentiation was associated with cognitive network reorganization seen during an episodic memory functional MRI paradigm, and correlated with inter-individual differences in task accuracy. Collectively, our findings showing a pattern of reduced microarchitectural differentiation between paralimbic regions and the remaining cortex provide a parsimonious explanation for functional network reorganization and cognitive dysfunction characteristic of TLE.

## Introduction

Temporal lobe epilepsy (TLE) is one of the most common pharmaco-resistant focal epilepsies ^1^. It is classically associated with variable degrees of mesiotemporal pathology ^2^ with many patients presenting with significant cognitive impairment, typically difficulties in learning and memory ^3-5^. Memory dysfunction impacts quality of life in patients with epilepsy ^6,7^, and affects several facets of daily living such as well-being, educational attainment, employment, and social functioning ^8^. These functional impairments emphasize the need to assess the neural processes underlying cognitive dysfunction in TLE. While early studies related memory impairment to structural changes in mesiotemporal regions ^9-13^, mounting evidence has shown distributed alterations in brain structure linking large-scale changes in grey matter volume ^14-16^, cortical morphology ^17-19^, and network properties ^20-25^ to cognitive dysfunction. These findings are in line with contemporary theories of memory, and its contributions to abstract thought more generally, which highlight the distributed nature of this function along with other higher-order cognitive abilities ^26-28^. These literatures underscore the importance of macroscale features of brain organization for our understanding of memory, and in particular their impairments in TLE. Our study set out to bring together these two lines of work with the aim of reconciling disease-related anomalies in neural architecture, distributed changes in memory circuit function, and resulting behavioural impairments in learning and memory, with emerging insights into the broad neural substrates that support memory-guided cognition.

Despite advances in our understanding of the structural correlates of memory dysfunction in TLE, the link between cognitive impairment and co-occurring changes in cortical microstructure remains elusive. Methodological and analytical advances in magnetic resonance imaging (MRI) acquisition and analysis have provided new means to investigate structure-function interactions in the human brain, enabling *in vivo* assessments of cortical microstructure and the organization of episodic memory networks in a patient-specific manner. In particular, emerging approaches in healthy populations applying unsupervised learning techniques to myelin-sensitive MRI contrasts, such as quantitative T1 (qT1) relaxometry, have paved the way for individualized investigations of intracortical microstructural organization in living humans and its association with cognitive topographies ^29-33^. In line with foundational *post mortem* neuroanatomical studies, these methods have outlined smooth and graded transitions in cortical laminar architecture running from unimodal sensory and motor cortices towards paralimbic circuits ^32,34^. By tracking region-to-region variations in intracortical lamination patterns, this *sensory-fugal gradient* of microstructural differentiation has been shown to recapitulate established models of cortico-cortical hierarchy ^30,32,35^. Episodic memory functions occupy a unique position in this hierarchy, co-localizing with transmodal functional systems such as paralimbic and default mode networks situated at the apex of the microstructural gradient ^26,32^. This topographic distance from sensory-motor systems at the interface of the external environment may be key for learning and remembering, as optimal memory functioning has been suggested to rely on the differentiation of neural activity of transmodal cortices from peripheral systems ^34,36-39^. Conversely, alterations in the microstructural substrate separating these systems may perturb this necessary dissociation, being a potential mechanism associated with memory dysfunction. Investigating topographic shifts of this gradient of cortical microstructure in TLE can, thus, identify macroscale changes in laminar organization in this condition, while offering a framework to explore cellular, molecular, and functional correlates of such changes.

Our goal was to assess large-scale changes in cortical microstructural differentiation in TLE and explore how these microstructural changes relate to memory dysfunction. We first compared cortex-wide *in vivo* microstructural gradients derived from high-resolution qT1 MRI between patients and controls. We observed reduced microstructural differentiation between paralimbic cortices and the remaining cortex in TLE, with epicentres in temporopolar and lateral prefrontal regions ipsilateral to the seizure focus. Stratifying our findings according to different cortical tissue types and harnessing *post mortem* histological and transcriptomics datasets, we demonstrated that disease-related microstructural gradient reorganization reflects topographical variations in cortical lamination and expression patterns of TLE-related risk genes.

Translating findings to cognitive architectures, we explored functional associations, with a focus on episodic memory networks. Specifically, we investigated cortical functional connectivity patterns of gradient reduction epicentres during the encoding phase of a functional MRI (fMRI) episodic memory task. We showed that regions of significant gradient perturbations harboured an atypical cognitive network architecture during memory fMRI, findings that were related to inter-individual differences in memory performance. Together, our findings define a novel framework to explore atypical structure-function coupling in TLE and the link between microstructural changes and cognitive abilities.

## Materials and methods

### Participants

#### a) Discovery sample

We studied 21 individuals with mesial TLE (9 women; mean age ± SD age 36.14 ± 11.59 years) referred to our centre for the investigation of pharmaco-resistant seizures. Patients underwent a research-dedicated 3T MRI between 2018 and 2021, which included qT1 MRI. Clinical history was obtained through interviews with patients and relatives. TLE diagnosis and lateralization of the seizure focus into left TLE (LTLE; n=15) and right TLE (RTLE; n=6) were determined by a comprehensive evaluation including detailed history, neurological examination, review of medical records, video-EEG recordings, and clinical MRI. Detailed clinical information including MRI-positive/negative status and suspected lesion type for each patient is provided in **TABLE S1**. Average age at seizure onset was 17.31±9.86 years (range=0.5-42 years) and epilepsy duration was 19.00±13.40 years (range=1-44 years). Six patients (28.57%) had a history of childhood febrile convulsions. At time of study, 2 patients had undergone a selective amygdalo-hippocampectomy, 4 had undergone cortico-amygdalo-hippocampectomy (including 1 patient with previous stereotactic radiofrequency thermocoagulation [RFTC] who were later referred for resective surgery due to seizure recurrence), 1 had undergone surgical resection of a ganglioglioma within the mesio-temporal lobe, and 3 had undergone RFTC of structures within the mesiotemporal lobe (no surgical resection). Seizure outcome was determined from Engel’s modified classification ^40^ with an average follow-up durations of 23.90±16.84 months. Since surgery, nine patients (90%) have been completely seizure-free (Engel-I); one remaining patient has recurrent seizures following RFTC and is currently awaiting surgery. Based on established histopathological criteria ^41^, four specimens showed hippocampal cell loss or gliosis. Histopathology of the resected hippocampus was unavailable in one of the temporal lobectomy cases, due to RFTC of the amygdala and hippocampus prior to resective surgery. History of severe traumatic brain injury and encephalitis was negative in all patients. The control group consisted of 35 individuals with no history of neurological or psychiatric conditions (HCs; 17 women; 34.51±9.27 years) who underwent the same imaging protocol as the TLE group ^42^. There were no significant differences in age (*t*=0.579; *p*=0.565) and sex (*Χ*^*2*^=0.172; *p*=0.678) between TLE and HC groups.

#### b) Replication sample

We assessed generalizability in an independent cohort of 22 TLE patients (13 women; 37.55±8.64 years) who underwent a research-dedicated 3T MRI in our institute between 2013 and 2016. A similar evaluation classified them as unilateral LTLE (n=9) or RTLE (n=13). Age at seizure onset was 12.45±10.80 years (range=1-36 years) and epilepsy duration was 25.00±11.44 years (range=2-39 years). Eleven patients (50%) had a history of childhood febrile convulsions. At time of study, 15 patients had undergone a selective amygdalo-hippocampectomy. Post-surgical follow-up durations varied between one and five years, with 11 patients becoming completely seizure-free (Engel-I) and four showing significant reduction in seizure frequency (Engel-II). All available specimens showed hippocampal cell loss or gliosis. History of severe traumatic brain injury and encephalitis was negative in all patients. Patients in the validation cohort were compared 18 HC (6 women; 34.78±7.72 years) who underwent the same imaging protocol. HCs reported a negative history of neurological or psychiatric illness. As in the discovery sample, there were no differences in age (*t*=1.057; *p*=0.297) and sex (*Χ*^*2*^=2.634; *p*=0.105) between patients and controls.

The Ethics Committee of the Montreal Neurological Institute and Hospital approved the study and written informed consent was obtained from all participants.

### MRI acquisition

#### a) Discovery sample

Scans were acquired at the McConnell Brain Imaging Centre (BIC) of the Montreal Neurological Institute and Hospital on a 3T Siemens Magnetom Prisma-Fit equipped with a 64-channel head coil. Participants underwent two T1-weighted (T1w) scans with identical parameters using a 3D magnetization-prepared rapid gradient-echo sequence (MPRAGE; 0.8 mm isovoxels, TR=2300 ms, TE=3.14 ms, TI=900 ms, flip angle=9°, FOV=256×256 mm^2^). T1w scans were inspected to ensure minimal head motion before undergoing further processing. qT1 relaxometry data were acquired using a 3D-MP2RAGE sequence (0.8 mm isovoxels, 240 sagittal slices, TR=5000 ms, TE=2.9 ms, TI_1_=940ms, T1_2_=2830 ms, flip angle_1_=4°, flip angle_2_=5°, FOV=256×256 mm^2^). We combined two inversion images for qT1 mapping to minimise sensitivity to B1 inhomogeneities and optimize reliability ^43,44^. Task and resting-state functional MRI (rsfMRI) scans were acquired using a 2D echo-planar imaging sequence (3.0mm isovoxels, matrix = 80×80, 48 slices oriented to AC-PC-30 degrees, TR=600ms, TE=30ms, flip angle=50°, multiband factor=6). The episodic memory task lasted ∼11 minutes (∼6 minutes for encoding and ∼5 minutes for retrieval). Stimuli were presented via a back-projection system. The rsfMRI scan lasted ∼7 minutes, and participants were instructed to fixate a cross displayed in the centre of the screen, to clear their mind, and not fall asleep.

#### b) Replication sample

Scans were also acquired at the BIC, using a 3T Siemens TrioTim with a 32-channel head coil. Participants underwent a 3D MPRAGE (1mm isovoxels, TR=2300 ms, TE=2.98 ms, TI=900 ms, flip angle=9°, FOV=256×256 mm^2^) and a 3D MP2RAGE sequence (1mm isovoxels, TR=5000 ms, TE=2.89 ms, TI_1_=940 ms, T1_2_=2830 ms, flip angle 1=4°, flip angle 2=5°, FOV=256×256 mm^2^) for T1w and qT1 imaging, respectively.

### Multimodal MRI processing and qT1 analysis

#### a) Multimodal preprocessing

Image processing leading to the extraction of cortical features and their registration to surface templates was performed via micapipe, an open multimodal MRI pipeline (http://github.com/MICA-MNI/micapipe/) ^45^. Subject-specific cortical surface models were generated from each native T1w scans using FreeSurfer 6.0 ^46-48^. Surface extractions were inspected and corrected for segmentation errors via placement of control points and manual edits. Native cortical features were registered to the Conte69 template surface (32,000 vertices per hemisphere) using workbench tools, and downsampled to 5,000 vertices/hemisphere. Task fMRI and rsfMRI images were pre-processed using AFNI ^49^ and FSL ^50^. The first five volumes of rsfMRI and task fMRI scans were discarded to ensure magnetic field saturation. Images were reoriented, as well as motion and distortion corrected. Motion correction was performed by registering all timepoint volumes to the mean volume, while distortion correction leveraged main phase and reverse phase field maps. Nuisance variable signal was removed using an ICA-FIX classifier ^51^. No global signal regression was performed. Volumetric timeseries were averaged for registration to native FreeSurfer space using boundary-based registration ^52^, and mapped to individual surfaces using trilinear interpolation. Cortical timeseries were mapped to the hemisphere-matched Conte69 template (with 32k surface vertices/hemisphere) using workbench tools ^53,54^ then spatially smoothed (Gaussian kernel, full width at half maximum [FWHM]=10mm). Surface- and template-mapped cortical timeseries were corrected for motion spikes using linear regression of motion outliers provided by FSL, and downsampled to retain 5k vertices per hemisphere.

#### b) Generation of microstructural gradients

We co-registered qT1 volumes to each participant’s native FreeSurfer space using boundary-based registration ^52^. We then generated 14 equivolumetric surfaces running between the pial and white matter boundaries to sample qT1 intensities across cortical depths in each participant, following validated methods (**FIGURE 1A**) ^32,55^. This procedure yielded distinct intensity profiles reflecting intracortical microstructural composition at each cortical vertex. Data sampled at the pial and white matter boundaries were discarded to mitigate partial volume effects. Intensity values at each depth were mapped to a common template surface, downsampled, and spatially smoothed across each surface independently (FWHM=3mm). Vertex-wise intensity profiles were cross-correlated using partial correlations controlling for the average cortex-wide intensity profile, and log-transformed. This resulted in microstructural profile covariance (MPC) matrices representing participant-specific similarity in myelin proxies across the cortex. We converted each participant’s MPC matrix to a normalized angle affinity matrix, and applied diffusion map embedding ^56^. This non-linear dimensionality reduction procedure identified eigenvectors (or *gradients*) describing main spatial axes of variance in inter-regional similarity of cortical microstructural profiles (**FIGURE 1B**). Procrustes analysis aligned subject-level gradients to a group-level template generated from the group-average MPC matrix of all TLE and HC participants in the discovery sample ^57,58^. Gradient analyses were performed using BrainSpace (http://github.com/MICA-MNI/BrainSpace), limiting the number of gradients to 10 and using default sparsity (keeping only the top 10% of MPC weights) and diffusion (α=0.5) parameters ^59^. Here, we focused on the principal gradient of microstructural similarity given its unique properties highlighting the differentiation of unimodal sensory and motor regions from transmodal cortices ^32^. Specifically, we tested how the topography of this gradient differed across participant groups.

**Figure 1.**
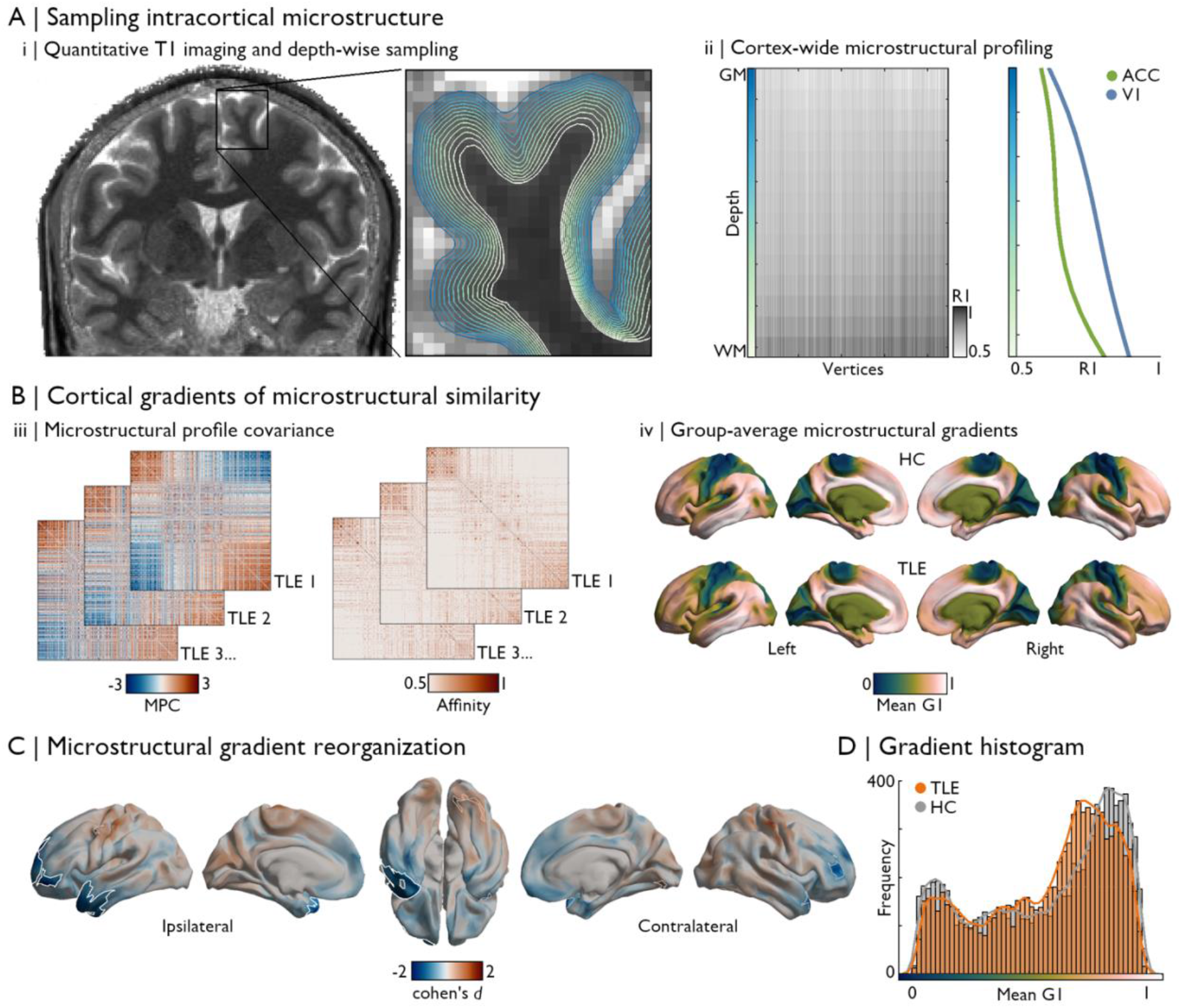
Microstructural gradient contractions in TLE. **(A)** Quantitative T1 imaging was used as a proxy of intracortical myeloarchitecture ^33^. Intensities were sampled at 14 cortical depths between the pial and white matter boundaries (*i*), yielding vertex-wise microstructural intensity profiles. T1 values were inversed (R1=1/T1) such that higher values reflect higher putative intracortical myelin content (*ii*), exemplified here by comparing profiles sampled in the primary visual cortex (V1, blue) and anterior cingulate cortex (ACC, green). **(B)** MPC matrices were constructed by cross-correlating vertex-wise intensity profiles using partial correlations controlling for the average cortex-wide profile, and normalized angle affinity matrices were generated from corresponding subject-level MPC matrices (*iii*). We applied dimensionality reduction techniques ^26,56^ to identify eigenvectors (gradients) describing main spatial axes in inter-regional similarity of cortical microstructural patterns ^32^. The average principal gradient (G1) is shown in (*iv*) for HC and TLE groups. **(C)** Surface-based linear models controlling for age and sex revealed significant differences in gradient scores between groups. Patients showed reductions in G1 scores in ipsilateral temporopolar and dorsolateral prefrontal cortices (*p*_FWE_<0.001). Significant clusters after multiple comparisons correction are outlined in white (*p*_FWE_<0.025), while trend-level effects are outlined in thinner grey clusters (*p*_FWE_<0.1). **(D)** Group-level histogram analysis confirmed that the distribution of G1 scores in TLE was compressed compared to controls, predominantly affecting paralimbic regions.

#### c) Statistical analysis

Left and right hemispheric data of individuals with TLE was sorted relative to the epileptogenic focus (*i*.*e*., into ipsi- and contralateral). To minimize confounds related to normal inter-hemispheric asymmetries, prior to sorting we normalized vertex-measures using z-transformations with respect to HCs ^60^. Gradient scores were compared between TLE and HCs using surface-based linear models implemented in BrainStat (http://github.com/MICA-MNI/BrainStat) ^61^. We controlled for age and sex, and corrected findings for family-wise errors (FWE) using random field theory (p_FWE_<0.025; cluster-defining threshold [CDT] = 0.01).

### Neural contextualization

#### a) Cytoarchitecture

Microstructural gradient findings in TLE were contextualized with respect to normative variations in cortical cytoarchitecture. The unthresholded effect size map (Cohen’s *d*) comparing age- and sex-corrected microstructural gradients in TLE and HCs was stratified according to cortical types defined by Von Economo and Koskinas ^62-64^. We also used the atlas proposed by Mesulam ^65^ defining four functional zones which follow variations in cortical laminar differentiation. We furthermore explored the association between microstructural gradient reorganization and histology-derived cortical cytoarchitectural differentiation. This analysis harnessed BigBrain, an ultra-high-resolution Merker-stained 3D reconstruction of a *post mortem* human brain in which staining intensities reflect neuronal size and density ^66^. Using analogous methods to those applied to derive *in vivo* microstructural gradients from qT1 imaging, gradients of cytoarchitectural differentiation were computed by sampling intracortical staining intensities in the BigBrain dataset. Histological staining intensity profiles and gradient maps are openly available as part of BigBrainWarp (http://github.com/caseypaquola/BigBrainWarp) ^67^. Two main axes of cytoarchitectural differentiation across the cortical mantle were explored, specifically sensory-fugal transitions differentiating sensory and motor cortices from paralimbic structures, and anterior-posterior gradients. Spearman rank correlations quantified the relationship between the topography of TLE-related microstructural gradient reorganization and these axes of cytoarchitectural differentiation. Statistical significance of correlations was determined using spin permutation tests (1,000 permutations) implemented in BrainSpace ^59^.

#### b) Disease-related transcriptomics

Microstructural gradient reorganization in TLE was contextualized against spatial transcriptomic signatures that are potentially relevant to epilepsy. For this purpose, we followed established procedures detailed in the ENIGMA toolbox (https://github.com/MICA-MNI/ENIGMA) ^64^, which aggregates preprocessed cortical gene expression maps from the Allen Human Brain Atlas (AHBA) dataset ^68-70^. Microarray samples were matched to parcels from the Schaefer-100 atlas ^71^ in each donor, and genes whose inter-donor correspondence fell below a pre-defined threshold (*r*<0.2) were discarded from further analysis. Gene lists provided in the ENIGMA toolbox include only those with biological prioritization in each epilepsy phenotype as identified in a previous genome-wide association study (GWAS) ^72^. We averaged expression maps of genes associated with focal hippocampal sclerosis (4 genes), all epilepsies (8 genes), generalized epilepsies (13 genes), juvenile myoclonic epilepsy (1 gene), childhood absence epilepsy (3 genes), and focal epilepsy (4 genes). The same parcellation as the gene expression data was applied to the unthresholded effect size map of microstructural gradient contractions to assess the spatial correlation between both datasets, with significance determined using spin permutation tests (1,000 permutations) as above. Included genes are listed in **TABLE S2**.

### Associations with memory circuits and function

To explore cognitive correlates of TLE-associated changes in microstructural differentiation, we assessed whether topography and magnitude of gradient changes related to cortical functional connectivity and performance during an episodic memory fMRI task. Experiment scripts and stimuli are openly available on GitHub (http://github.com/MICA-MNI/micaopen/, see *task-fMRI*).

#### a) Task design and analysis sample

The memory task consisted of two phases. First, participants completed an encoding phase in which they were instructed to memorize 56 image pairs consisting of realistic depictions of common objects. A subset of pairs was shown twice, for a total of 84 trials. Trial duration was 2 seconds and were separated by a randomly jittered 1.5 to 3.5-second interval. Approximately 12 minutes after the encoding phase, participants were asked to match an image shown at the top of the screen with its paired image from the encoding phase in a 3-alternative forced choice design. After excluding participants with incomplete data (n=4) and below chance-level performance (n=4), a total of 48 participants were retained for this analysis (16 TLE and 32 HC).

#### b) Task-based and resting-state functional connectivity

To study how regions showing atypical microstructural differentiation were embedded within functional memory networks, we selected seed vertices corresponding to peak regions in each cluster of significant gradient reductions (*i*.*e*., one peak in the temporopolar cluster and one peak in the lateral prefrontal cluster). Seed timeseries were correlated with the timeseries of all other vertices across the cortex. Correlation coefficients underwent Fisher’s *r*-to-z transformation, and connectivity maps were compared across patient and controls groups using surface-based linear models accounting for age and sex. This analysis was performed using functional timeseries collected during the encoding phase. To assess the specificity of uncovered connectivity alterations during memory encoding, we repeated this analysis using timeseries extracted from a resting-state fMRI paradigm acquired during the same session (and processed using the same pipelines).

#### c) Post hoc analysis of recall accuracy

*Post hoc* analyses exploring the association between gradient changes and behavioural task performance were restricted to clusters of significant gradient reductions. Age- and sex-corrected z-scores were extracted from individual gradient maps at each seed described in *b)*, and correlated with individual recall performance. Data from all TLE and HC participants were pooled for this analysis.

## Results

### Robust and specific microstructural gradient alterations in TLE

Cortex-wide intracortical microstructural profiles were generated for each participant. These profiles consisted of qT1 image intensities sampled along 14 equivolumetric surfaces running between the pial and white matter boundaries (**FIGURE 1A**). Cross-correlating vertex-wise intensity profiles resulted in subject-specific matrices representing similarity in myelin proxies across the cortex. We estimated eigenvectors describing spatial gradients of cortex-wide microstructural variations (**FIGURE 1B**). Using this gradient in case-control comparisons can uncover different patterns of alterations in patients relative to controls. For instance, an expansion of the gradient in the TLE group (or scores shifting further away from the gradient midpoint) would suggest that local perturbations in cortical microstructure are further increasing inter-areal differentiation. However, a contraction of the gradient (or scores shifting towards the gradient midpoint) would indicate overall less pronounced microstructural differentiation in affected regions of patients.

As in previous *post mortem* histology and *in vivo* MRI findings in healthy individuals, the average principal gradient of cortical microstructure in TLE and HC groups differentiated primary sensory and motor regions from paralimbic cortices ^30,32^. Compared to controls, individuals with TLE showed marked reductions in microstructural gradient scores in anterior temporal (temporopolar, lateral temporal; *p*_FWE_<0.001) and prefrontal regions (frontopolar, dorsolateral prefrontal; *p*_FWE_<0.001) ipsilateral to the seizure focus (**FIGURE 1C**). More subtle gradient reductions were also seen in homologous contralateral regions (contralateral temporal pole: *p*_FWE_<0.05; contralateral ventrolateral prefrontal cortex: *p*_FWE_<0.05). Increases in gradient scores in TLE compared to controls were concentrated in unimodal sensory and motor cortices, and were strongest in contralateral inferior occipito-temporal regions (*p*_FWE_<0.05) and trended in the ipsilateral precentral gyrus (*p*_FWE_=0.07). In line with surface-based analyses, the histogram of group-average gradient values showed overall gradient reductions in patients with TLE relative to controls (Wilcoxon rank sum test: *z*=-4.147; *p*<0.001). The TLE group showed prominent decreases in the paralimbic extreme of the microstructural gradient, but only subtle increases in the sensory-motor anchor (**FIGURE 1D**). *Post hoc* analyses focusing on significant clusters of gradient reductions demonstrated significantly higher microstructural profile similarity of temporopolar and dorsolateral prefrontal regions to unimodal sensory and motor cortices, as well as reduced similarity to paralimbic and transmodal cortices in the TLE group (**FIGURE S1**). These results are compatible with findings observed at the gradient-level indicating large-scale patterns of microstructural dedifferentiation of the cortex in TLE. These findings highlight a contracted myeloarchitectonic gradient in TLE, with reduced differentiation of paralimbic and transmodal regions from the rest of the cortex, particularly affecting ipsilateral temporopolar and dorsolateral prefrontal cortices.

Robustness of microstructural gradient contractions was assessed with several additional analyses (see **SUPPLEMENTARY MATERIALS for details**). First, we found gradient reorganizations to be consistent across individuals, with >80% of patients showing at least moderate reductions in gradient scores (*z*<-1) in either significant cluster of findings (**FIGURE S2)**. Lower z-scores in either significant cluster (averaged within each cluster) could also be seen in the ipsilateral vs. contralateral hemisphere in >90% of patients (**FIGURE S2)**. In addition, investigations of secondary eigenvectors explaining less variance than the principal microstructural gradient confirmed consistent changes in ipsilateral anterior temporal regions (**FIGURE S3**). Findings were similar when studying microstructural differentiation using an alternative approach *i*.*e*., depth-dependent statistical moments quantifying the shape of intracortical intensity profiles (**SUPPLEMENTARY METHODS and FIGURES S4, S5**). Moreover, results were robust when locally controlling for cortical thinning and pericortical blurring, two common imaging findings previously reported in TLE ^73-75^, suggesting intralaminar specificity (**SUPPLEMENTARY METHODS and FIGURE S6**). Finally, microstructural gradient contractions were replicable in a distinct cohort of participants who showed prominent gradient contractions across the ipsilateral temporal lobe, with significant reductions in anterior mesial and inferior temporal regions (**FIGURE S7**).

### Microarchitectural underpinnings of gradient reorganizations

Cytoarchitectural stratification of microstructural gradient contractions supported the interpretation that disease-related changes in the principal microstructural gradient reflected a large-scale dedifferentiation of cortical microstructure. Grouping gradient effect sizes according to atlases of cortical types and functional zones confirmed that gradient reductions were strongest in paralimbic and dysgranular cortices, while gradient increases in TLE tended to affect primary areas with strong laminar differentiation (**FIGURE 2A**). Gradient changes across the cortex furthermore spatially correlated with sensory-fugal cytoarchitectural differentiation derived from 3D histological data ^66,67^ (r=-0.527, p_spin_=0.001). Here, gradient reductions in TLE co-localized with paralimbic areas, while gradient increases overlapped with primary sensory-motor regions (**FIGURE 2A**). However, cortex-wide correlations with the anterior-posterior axis of cytoarchitectural variation were not significant (r=-0.278, p_spin_=0.252). These findings demonstrate that macroscale perturbations in cortical microstructure in TLE follow the overall axis of cortical cytoarchitectural differentiation mapped from *post mortem* histology.

**Figure 2.**
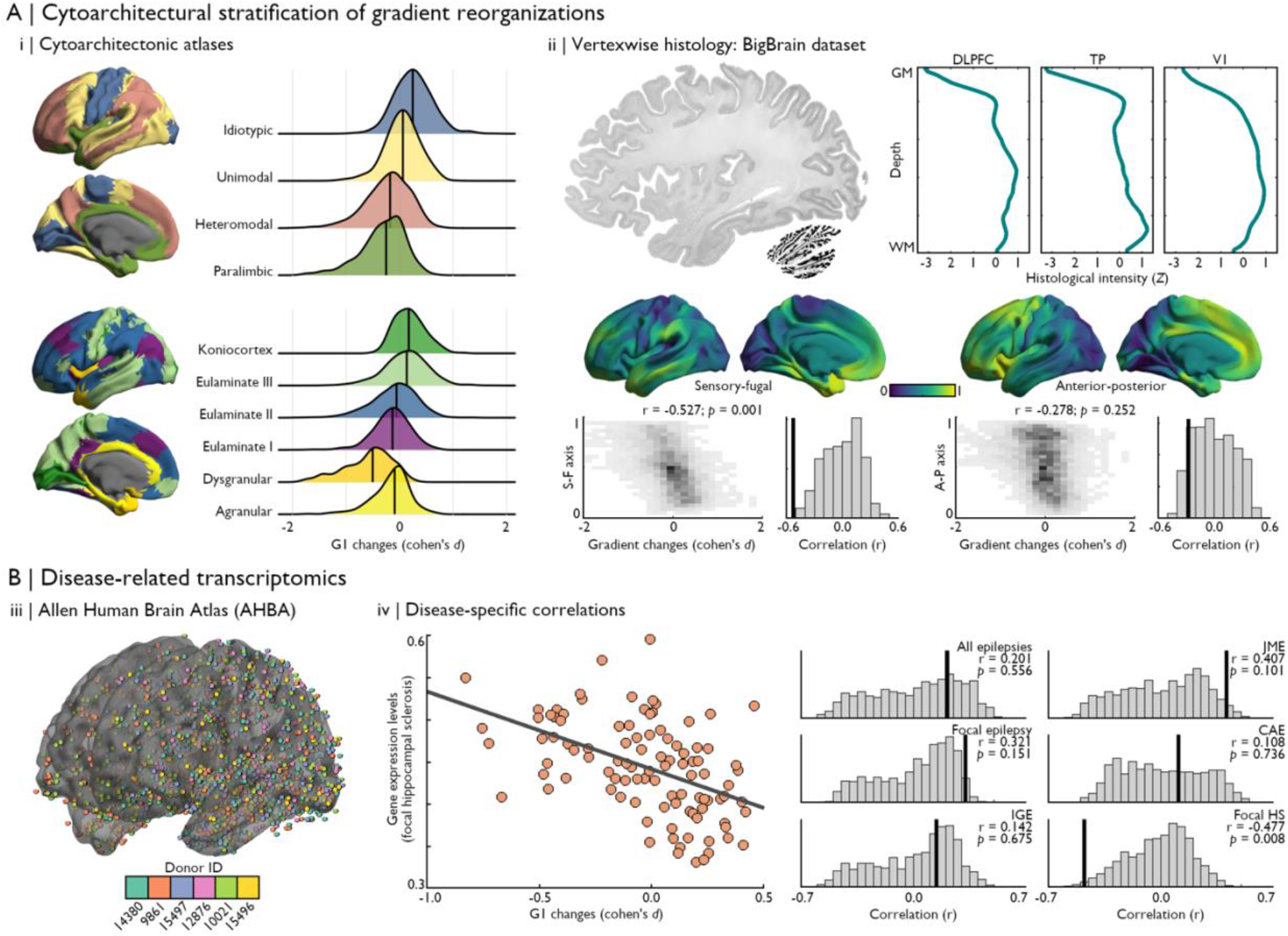
Cytoarchitectural and transcriptomic contextualization of microstructural gradient contractions. **(A)** *(i)* In reference to atlases of cortical types and functional zones, we found that gradient 1 (G1) score reductions were strongest in paralimbic and dysgranular cortices, while areas of TLE-related G1 increases co-localized with highly differentiated idiotypic areas as well as eulaminate III and koniocortical regions. *(ii)* 3D histological data from the BigBrain dataset ^66^ (left) was pre-processed and made available on various surface templates in the BigBrainWarp toolbox ^67^. Selected examples of vertex-wise histological profiles in the dorsolateral prefrontal cortex (DLPFC), temporal pole (TP), and primary visual cortex (V1) highlight heterogeneous cytoarchitectural characteristics across regions. Main axes of variations in cytoarchitectural similarity across the cortex were seen along sensory-fugal and anterior-posterior directions. The topography of microstructural gradient changes in TLE was significantly correlated with the sensory-fugal pattern of cytoarchitectural differentiation (*r*=-0.527; *p*_spin_=0.001), but not with the anterior-posterior axis (*r*=-0.278; *p*_spin_=0.252). **(B)** *(iii)* Leveraging gene expression data of the AHBA ^69^, we assessed the specificity of microstructural gradient alterations to TLE-related disease processes. *(iv)* Pre-processed gene expression and gene lists ^72^ available in the ENIGMA toolbox ^64^ were used in disease-related transcriptomic analyses. The topography of TLE-related microstructural gradient contractions followed the average expression patterns of genes related to hippocampal sclerosis (*r*=-0.477, *p*_spin_=0.008). No significant correlations were found with other epilepsy type-related gene expression patterns. Statistical significance of correlations with histological and transcriptomic features was determined using non-parametric spin permutation testing (1,000 permutations).

Next, we leveraged disease-related transcriptomics analyses to contextualize macroscale changes in microstructural differentiation with respect to gene expression patterns associated with different epilepsy syndromes. We mapped the spatial expression profiles of disease-related risk genes (obtained from recent GWAS ^72^) using data from the AHBA (for a similar approach, see ^45^). Average expression maps of genes associated with different epilepsy types (*i*.*e*., epilepsy-related hippocampal sclerosis, focal epilepsy, juvenile myoclonic epilepsy, childhood absence epilepsy, generalized epilepsy, all epilepsies) were first correlated with the parcellated effect size map of TLE-related gradient reorganizations. We observed a correlation between patterns of microstructural gradient changes and risk gene expression levels of hippocampal sclerosis even after controlling for spatial autocorrelation (*r*=-0.477, *p*_spin_=0.006). Notably, correlations with other epilepsy-related gene sets were non-significant (*focal epilepsy*: *r*=0.321, *p*_spin_=0.151; *juvenile myoclonic epilepsy*: *r*=0.407, *p*_spin_=0.101; *childhood absence epilepsy*: *r*=0.108, *p*_spin_=0.736; *idiopathic generelized epilepsy*: *r*=0.142, *p*_spin_=0.675; *all epilepsies*: *r*=0.201, *p*_spin_=0.556; **FIGURE 2B**). Syndrome-specific gene expression maps are displayed in **FIGURE S8**. These results highlight the relative specificity of macroscale perturbations in cortical microstructural differentiation to regions with higher vulnerability to TLE-related genetic processes.

### Associations to cognitive network reorganization and impairment

After establishing the topography of TLE-related changes in cortical microstructural gradients, we studied how identified regions were embedded within functional memory networks. Episodic memory fMRI involved image-pair learning followed by retrieval testing in a three-alternative forced choice design across 56 trials (**FIGURE 3A**). Encoding task timeseries were extracted from two seed vertices centered on peak regions in clusters of gradient reductions, specifically in temporopolar and lateral prefrontal regions ipsilateral to the seizure focus (**FIGURE 3B**). We observed reduced connectivity between both seeds, with stronger effects observed in the hemisphere ipsilateral to the seizure focus (**FIGURE 3B)**. Comparing connectivity patterns of each seed across groups using surface-based linear models were consistent with these results. Significant clusters of connectivity reductions during memory encoding were found between the temporopolar seed and orbitofrontal, ventromedial, and ventrolateral prefrontal regions. Relatedly, significant connectivity reductions were found between the lateral prefrontal seed and lateral and inferior temporal cortices. Interestingly, the topography of seed-based connectivity reductions during encoding did not generalize to intrinsic functional connectivity profiles of the same seeds computed from a resting-state scan (**FIGURE S9**). Moreover, microstructural gradient alterations in these regions were associated with task performance. After controlling individual recall performance and gradient scores for age and sex, we found significant correlations between episodic memory function and microstructural gradient reorganizations in temporopolar regions (*r*=0.406, *p*=0.004), but not lateral prefrontal regions (*r*=0.111, *p*=0.457; **FIGURE 3C**). These results suggest that microstructural dedifferentiation in TLE, and particularly changes involving ipsilateral temporal structures, are associated with dysfunction of higher-order cognitive processes such as episodic memory supported by fronto-temporal circuits.

**Figure 3.**
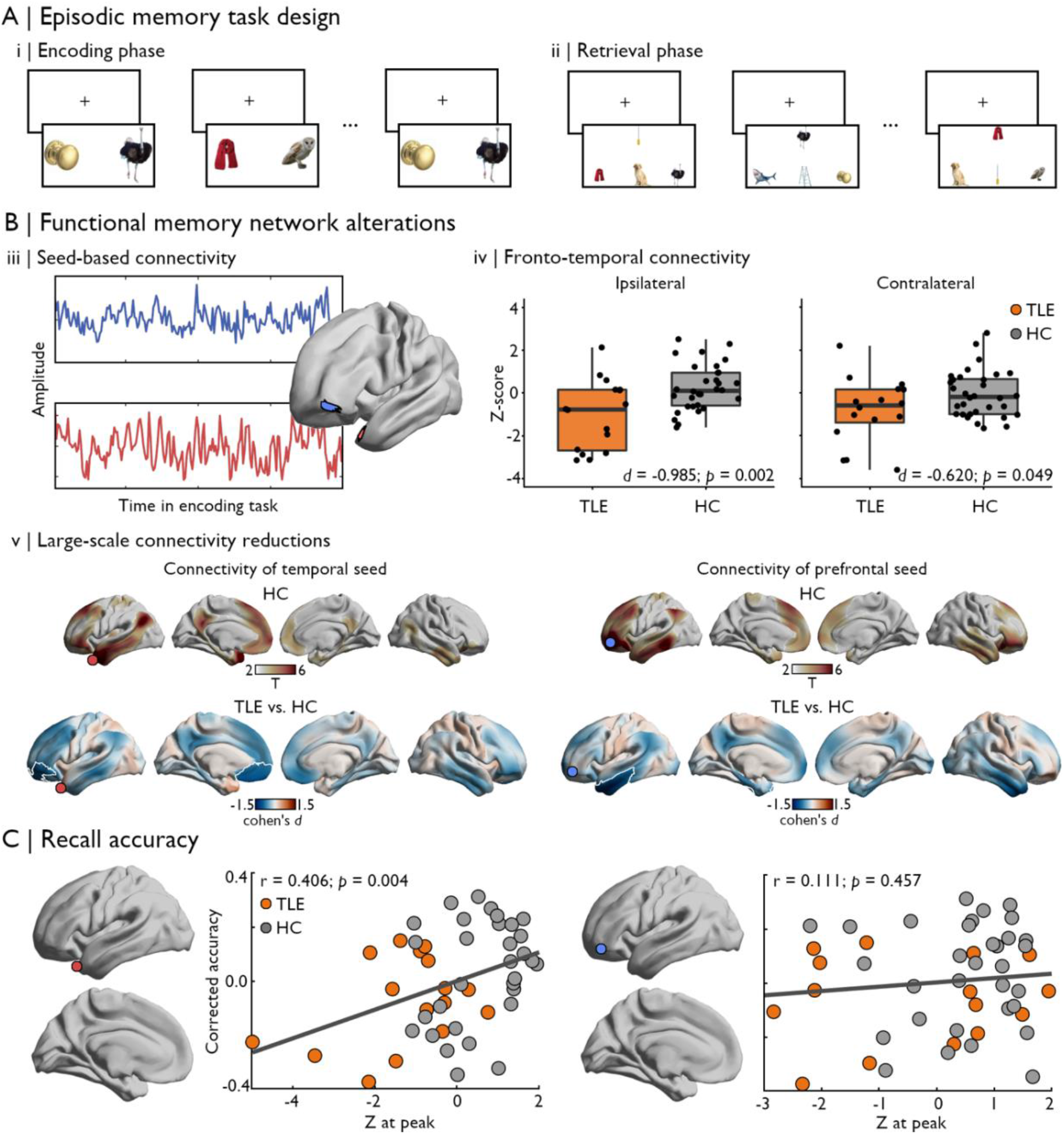
Microstructural gradient changes are related to episodic memory network configurations. **(A)** Participants completed an episodic memory task involving encoding and recall of 56 distinct image pairs, with half of pairs shown twice during the encoding phase. The task involved distinct encoding *(i)* and retrieval phases *(ii)*, separated by a 12-minute delay. **(B)** Seed-based functional connectivity analysis during the encoding task centered on peak regions in clusters of gradient reductions *(iii)* showed significantly reduced connectivity between temporal and prefrontal regions, with more pronounced effects ipsilateral to the seizure focus *(iv)*. These findings were consistent with cortex-wide connectivity changes of each seed, which also revealed more extended connectivity reductions in TLE involving default mode and limbic subregions *(v)*. In HCs (top), regions with significant functional connections to temporal (left) and prefrontal (right) seeds are depicted in coloured regions on the cortical surface (*p*_FWE_<0.05). The bottom panel illustrates TLE-related changes in functional connectivity during memory encoding, with significant clusters of connectivity reductions relative to HCs outlined in white (*p*_FWE_<0.05) over the unthresholded effect size map. The location of each seed region is indicated by a coloured marker. **(C)** Microstructural gradient changes in temporopolar (left), but not prefrontal regions (right), were significantly correlated with recall accuracy, with individuals showing stronger gradient contractions also scoring lower in the retrieval phase of this episodic memory task.

## DISCUSSION

An emerging literature emphasizes the importance of brain-wide topography in understanding structure-function links and cognitive architectures in both health and disease. Here, we capitalized on a high-resolution neuroimaging dataset to explore cortical microstructural differentiation in temporal lobe epilepsy (TLE), one of the most common pharmaco-resistant epilepsies in adults, and to clarify its relationship with memory dysfunction commonly observed in this condition. We used a novel approach to conceptualize focal microstructural alterations in TLE within their broader architectural context using large-scale gradients of cortical myeloarchitecture. We found a significantly contracted *sensory-fugal* gradient in patients, reflecting large-scale dedifferentiation of cortical microstructure between paralimbic cortices and the remaining cortex, with epicentres located in temporopolar and dorsolateral prefrontal regions ipsilateral to the seizure focus. Microstructural gradient reorganization observed in patients reflected topographical variations in cortical lamination patterns and critically followed gene expression profiles associated with hippocampal sclerosis, emphasizing the relative specificity of our findings to TLE-related processes. Lastly, we showed that regions of strongest microstructural gradient contractions captured perturbed fronto-temporal connectivity during encoding of new material and were significantly correlated with recall performance from the same task. Collectively, our work demonstrates the association between disease-related gradient alterations and cortical cytoarchitecture, gene expression, memory network organization, as well as memory performance, while supporting the pivotal role of this sensory-fugal gradient for memory dysfunction in this cohort. Indeed, TLE appears to change associations between the microstructure of paralimbic regions and the remaining cortex in a way that *(a)* is consistent with what is already known about gene expression in this condition, and *(b)* recapitulates topographical themes thought to be important in emerging perspectives on how memory can guide behaviour.

Hippocampal sclerosis and microstructural changes within neighboring mesiotemporal structures are a core feature of TLE ^41,76-78^. Recent advances in high-resolution MRI have facilitated investigations of these alterations, opening the way to detailed *in vivo* characterizations of temporal and extra-temporal microstructural pathology. Previous studies harnessing different microstructurally-sensitive MRI contrasts have reported robust signal changes within the mesiotemporal disease epicentre. In line with histopathological examinations of extratemporal neocortical tissue ^79^, these investigations have also shown more widespread alterations affecting paralimbic and transmodal association networks, including temporopolar, lateral temporal, orbitofrontal, and prefrontal regions ^23,80,81^. Moreover, recent work using qT1 imaging described the depth-specific nature of these changes, reporting stronger and more spatially distributed signal changes in more superficial layers ^80^. These findings are compatible with previous histological reports on atypical cortical layering occurring in conjunction with abnormal horizontal fibres and clusters of small neurons in upper cortical layers in patients with TLE ^82-85^. Our approach builds on these findings by considering local alterations in laminar architecture within their broader microstructural context, captured by subject-specific depictions of microstructural similarity patterns across the cortex. The macroscale heterogeneity of cortical microstructure is a key feature of brain organization. Neuronal migration guided by signalling molecules is an important contributor to the patterning of laminar subtypes across the adult neocortex. This migration process terminates according to different timelines across cytoarchitectonic areas ^86^, and defects in this prenatal process are associated with several epilepsy types. Notably, dysfunction of Reelin signalling pathways, a core component of neuronal migration, has been associated with hippocampal sclerosis but may also affect laminar patterning of more extended neocortical sites ^87-90^. Regional laminar architecture and inter-areal similarity of laminar properties are closely linked to cortico-cortical connectivity patterns, as regions with similar laminar organization are more likely to share strong connections ^91-93^. Therefore, large-scale imbalances in the microstructural differentiation of the cortex have the potential to shift the balance of integration and segregation within brain networks. Structural and functional network topology are known to be altered in TLE, and frequently involve temporo-limbic and higher-order association cortices ^94-98^, co-localizing with significant clusters of findings identified in the present study. We speculate that distributed and heterogeneous anomalies in regional laminar architecture observed in TLE may contribute to reconfigurations of cortical networks frequently observed in this condition. In turn, these perturbations may be associated with the generation and propagation of seizures, but also to other phenotypic aspects of this condition, such as cognitive dysfunction.

Our understanding of the human brain has increasingly benefitted from ‘multiscale’ approaches integrating microarchitectural features with macroscale principles of brain organization ^68,93,99-101^, a perspective that is also influential in contemporary thinking in how memory guides our thoughts and behaviour. To better contextualize TLE-related changes in sensory-fugal gradient topography, our study capitalized on measures of cytoarchitectural laminar architecture. We found that these changes followed variations in cortical laminar differentiation patterns, confirming that the altered microstructural embedding space identified from in vivo qT1 imaging reflected increased proximity of regions separated by differences in their laminar structure. Notably, strongest effects were captured by reductions of gradient scores within paralimbic/dysgranular cortical subtypes. Disposed in a ring-like formation at the base of the cortex, limbic and paralimbic cortices are characterized by little differentiation across layers ^34,91,102^. Several cytoarchitectural and molecular properties of limbic areas are thought to facilitate their greater plasticity relative to regions with a more defined laminar structure ^103^. For instance, different growth-associated proteins are predominantly expressed in limbic and association cortices during adulthood relative to primary sensory and motor cortices ^91,104,105^. However, this increased plasticity may render limbic areas more vulnerable to pathology ^91^ and create a favourable context for the emergence of more widespread microstructural anomalies and associated epileptogenic networks within these regions ^106^. For instance, animal models of TLE have shown that increased plasticity (*e*.*g*., indexed by the increased expression of molecules such as GAP-43) may contribute to the development of epileptogenic networks and the early stages of mossy fibre sprouting ^107-109^. Several control analyses could furthermore demonstrate the specificity of our findings for intracortical processes, as results were stable when independently controlling for subject-level measures of cortical thickness and peri-cortical interface contrast, two morphological and intensity features that may be atypical in TLE. In contrast, TLE-associated reorganization of the sensory-fugal gradient strongly co-localized with changes in microstructural intensity profiles sampled within the cortical sheet. Quantitative profiling of local laminar architecture using statistical moments is a well-established approach for areal boundary definition in histological studies ^110,111^ and has only recently been translated to *in vivo* data ^29,31^. Although our study demonstrates the potential of this method in characterizing widespread microstructural changes associated with TLE, precise investigations of underlying cytoarchitectural changes should be further explored using patient-specific quantitative histopathological data ^112,113^.

Microstructural gradient contractions in TLE were also found to co-localize with disease-specific molecular features. Brain microstructure and gene expression are intricately linked: Distinct transcriptomic signatures are associated with different neural cell types ^114-116^, cortical laminar architectures ^116-118^, and laminar patterning of cortico-cortical projections ^119^. However, the transcriptomic mechanisms supporting macroscale brain organization, and particularly how atypical molecular function ultimately contributes to clinical phenotypes, remain relatively unclear. By combining an openly available microarray transcriptomic dataset with results of a recent epilepsy GWAS, we show that sensory-fugal gradient contractions seen in TLE followed spatial expression patterns of risk genes associated with hippocampal sclerosis. Indeed, microstructural gradient reductions seen in temporo-limbic and higher-order association cortices co-localized with areas of elevated expression of TLE risk genes, while correlations with risk gene expression associated with other epilepsy syndromes were not significant. Similarly, a recent study identified associations of structural covariance network reconfigurations in TLE with disease-related transcriptomic patterns ^70^. Although these results offer potential insights into the transcriptomic basis of macroscale pathophysiological processes associated with TLE, the combination of normative gene expression patterns obtained from the AHBA dataset and GWAS findings is hindered by several limitations ^70^. Firstly, while AHBA offers excellent cortical coverage, data are derived from only six healthy donor brains with predominant sampling in the left hemisphere ^69^. Moreover, the AHBA dataset relies on microarray transcriptomics, which may offer reduced sensitivity and specificity to map gene expression compared to more elaborate and costly methods such as RNA-Seq. Finally, our understanding of epilepsy risk genes identified by GWAS may also evolve as more patients are enrolled in these studies, given that GWAS will likely identify more relevant genes with increasing sample sizes ^120^. Despite these unique methodological challenges, our results indicate that macroscale perturbations in sensory-fugal organization in TLE may be rooted in disease-specific genetic processes.

Previous studies have linked the structural disease substrate of TLE to cognitive impairment ^19,21^, including episodic memory deficits classically associated with the condition ^3^. Notably, measures looking beyond regional changes and considering macroscale alterations in brain structure and network organization can accurately capture profiles of neuropsychological impairment in TLE ^19-22,121^. In line with this approach, we investigated disease-related changes in functional connectivity of areas showing maximal microstructural gradient contractions during relational episodic memory encoding. Our findings revealed that regions with abnormal microstructure in TLE are also functionally disconnected to temporo-limbic and default mode subregions. Indeed, anterior temporal regions showed significantly reduced connectivity to ventrolateral and orbitofrontal regions, while lateral prefrontal regions were less connected to lateral and inferior temporal regions relative to controls during this task. Frontotemporal circuits involve crucial nodes for episodic memory function in humans, and perturbations in these networks may impair encoding and retrieval of newly learned material ^122,123^. Several studies have investigated the reorganization of functional memory networks in TLE, consistently showing increased lateralization of memory function to the contralesional hemisphere ^124-127^. In line with this observation, we found that encoding network connectivity reductions followed the lateralization of microstructural gradient alterations, with stronger ipsilesional connectivity reductions in frontotemporal circuits. These findings are consistent with a recent study showing perturbed functional connectivity between the mesio-temporal lobes and more distant frontal nodes supporting episodic memory processes ^128^. Increased recruitment of extra-temporal structures during memory encoding has also been demonstrated in TLE, notably involving regions of the dorsolateral prefrontal cortex ^124,129^. Greater activation of several extra-temporal paralimbic structures in TLE, such as insular, anterior cingulate, and orbitofrontal cortices have been linked with better verbal learning performance ^124^. This greater recruitment of extratemporal and contralateral temporal regions has been proposed to reflect a potentially compensatory, yet suboptimal, mechanism to maximize encoding of new material that leverages the extended network supporting episodic memory function ^130^. Our results suggest that underlying microstructural damage in TLE may be associated with relative disconnection of crucial nodes within this network, possibly underpinning the inefficacy of this compensatory mechanism. Our behavioural analysis showed that more marked microstructural gradient reductions in temporal regions were significantly correlated with worse recall performance. Collectively, these findings offer novel evidence of perturbed structure-function coupling in TLE, that not only affects the mesio-temporal disease epicentre, but also extends to frontotemporal systems supporting memory encoding. Moving forward, this work provides an important analytic approach to leverage TLE as a disease model for how remembering can fail, and for understanding how memory can guide cognition in a more general manner.

Several control analyses supported specificity and robustness of our main findings. First, significant group-level patterns of microstructural gradient contractions were reproducible in most patients. Patient-specific asymmetry in microstructural gradient scores were furthermore consistent with lateralization of the seizure focus in over 90% of cases. These findings support the potential clinical utility of considering sensory-fugal gradient reorganizations for diagnostics; replication in larger cohorts and using different microstructure-sensitive imaging contrasts are, however, warranted. In this regard, we replicated the main patterns of findings in an independent cohort of TLE patients and healthy controls who underwent similar imaging on a different MRI scanner, and showed significant gradient contractions in the temporal lobe, with stronger effects observed ipsilateral to the seizure focus. However, the prefrontal gradient changes that were observed in the discovery cohort did not reach significance after multiple comparisons correction in the validation dataset. TLE diagnosis was supported by favourable post-surgical outcomes in most operated cases in both cohorts. Despite careful consideration of available clinical data, the slightly higher proportion of unoperated cases and shorter follow-up times in our discovery cohort may indicate potential differences in the spatial extent and severity of pathology across samples.

In summary, the present work highlights the potential of moving beyond regional analyses to study cortical microstructural anomalies in TLE. Microstructural changes in this condition exist within a broader architectural context and imply a cortex-wide dedifferentiation. This work thus centres well-documented local microstructural alterations within our contemporary understanding of TLE as a network disorder. Findings aligned with important axes of cortical laminar differentiation and with the expression patterns of genes associated with hippocampal sclerosis. Finally, microstructural gradient changes also related to memory performance and reorganization of associated functional networks. Collectively, these results provide a novel way to conceptualize microstructural features of TLE, while opening new avenues to investigate their contribution to clinical presentations and decision making.

## ACKNOWLEDGEMENTS

The authors would like to thank all patients and control participants who took part in this study.

## COMPETING INTERESTS

The authors report no competing interests.

## FUNDING

J.R. was funded by a fellowship from the Canadian Institutes of Health Research (CIHR). S.L. was supported by a CIHR doctoral scholarship and the Ann and Richard Sievers Award in Neuroscience. R.R.C. received funding from the Fonds de la Recherche du Québec – Santé (FRQS). BP was supported by the National Research Foundation of Korea (NRF-2021R1F1A1052303; NRF-2022R1A5A7033499), Institute for Information and Communications Technology Planning and Evaluation (IITP) funded by the Korea Government (MSIT) (No. 2022-0-00448, Deep Total Recall: Continual Learning for Human-Like Recall of Artificial Neural Networks; No. 2020-0-01389, Artificial Intelligence Convergence Research Center (Inha University); No. RS-2022-00155915, Artificial Intelligence Convergence Innovation Human Resources Development (Inha University); No. 2021-0-02068, Artificial Intelligence Innovation Hub), and Institute for Basic Science (IBS-R015-D1). L.C. acknowledges support from a Berkeley Fellowship jointly awarded by UCL and Gonville and Caius College, Cambridge, and by Brain Research UK (award 14181). A.B. and N.B. were supported by FRQS and CIHR. B.F. was supported by a salary award of the FRQS (Chercheur-boursier Senior) and acknowledges support from the National Science and Engineering Research Council of Canada (NSERC) and CIHR. B.C.B. acknowledges support from NSERC, CIHR, SickKids Foundation, Azrieli Center for Autism Research (ACAR-TACC), BrainCanada, Future Leaders Research Grant, Helmholtz International BigBrain Analytics and Learning Laboratory (HIBALL), FRQS, and the Canada Research Chairs program.

## Notes

### Competing Interest Statement

The authors have declared no competing interest.

## REFERENCES

1. Wieser H-G. ILAE Commission Report. Mesial temporal lobe epilepsy with hippocampal sclerosis. Epilepsia. 2004;45(6):695–714.

2. Blumcke I, Spreafico R, Haaker G, et al. Histopathological findings in brain tissue obtained during epilepsy surgery. New England Journal of Medicine. 2017;377(17):1648–1656.

3. Bell B, Lin JJ, Seidenberg M, Hermann B. The neurobiology of cognitive disorders in temporal lobe epilepsy. Nature Reviews Neurology. 2011;7(3):154–164.

4. Hermann B, Seidenberg M, Lee E-J, Chan F, Rutecki P. Cognitive phenotypes in temporal lobe epilepsy. Journal of the International Neuropsychological Society. 2007;13(1):12–20.

5. Saling MM. Verbal memory in mesial temporal lobe epilepsy: beyond material specificity. Brain. 2009;132(3):570–582. doi:10.1093/brain/awp012

6. Perrine K, Hermann BP, Meador KJ, et al. The relationship of neuropsychological functioning to quality of life in epilepsy. Archives of Neurology. 1995;52(10):997–1003.

7. Giovagnoli AR, Avanzini G. Quality of life and memory performance in patients with temporal lobe epilepsy. Acta Neurologica Scandinavica. 2000;101(5):295–300. doi:https://doi.org/10.1034/j.1600-0404.2000.90257a.x

8. Baker GA, Taylor J, Hermann B. How can cognitive status predispose to psychological impairment? Epilepsy & Behavior. 2009/06/01/ 2009;15(2, Supplement 1):S31–S35. doi:https://doi.org/10.1016/j.yebeh.2009.03.021

9. Rausch R, Babb TL. Hippocampal Neuron Loss and Memory Scores Before and After Temporal Lobe Surgery for Epilepsy. Archives of Neurology. 1993;50(8):812–817. doi:10.1001/archneur.1993.00540080023008

10. Lencz T, McCarthy G, Bronen RA, et al. Quantitative magnetic resonance imaging in temporal lobe epilepsy: Relationship to neuropathology and neuropsychological function. Annals of Neurology. 1992;31(6):629–637. doi:https://doi.org/10.1002/ana.410310610

11. Kilpatrick C, Murrie V, Cook M, Andrewes D, Desmond P, Hopper J. Degree of left hippocampal atrophy correlates with severity of neuropsychological deficits. Seizure. 1997;6(3):213–218.

12. Baxendale S, Van Paesschen W, Thompson P, et al. The relationship between quantitative MRI and neuropsychological functioning in temporal lobe epilepsy. Epilepsia. 1998;39(2):158–166.

13. Reminger SL, Kaszniak AW, Labiner DM, et al. Bilateral hippocampal volume predicts verbal memory function in temporal lobe epilepsy. Epilepsy & Behavior. 2004;5(5):687–695.

14. Bonilha L, Alessio A, Rorden C, et al. Extrahippocampal gray matter atrophy and memory impairment in patients with medial temporal lobe epilepsy. Human Brain Mapping. 2007;28(12):1376–1390. doi:https://doi.org/10.1002/hbm.20373

15. Hermann B, Seidenberg M, Bell B, Rutecki P, Wendt G, Magnotta V. Extratemporal quantitative MR volumetrics and neuropsychological status in temporal lobe epilepsy. Journal of the International Neuropsychological Society. 2003;9(3):353–362.

16. Focke NK, Thompson PJ, Duncan JS. Correlation of cognitive functions with voxel-based morphometry in patients with hippocampal sclerosis. Epilepsy & Behavior. 2008;12(3):472–476.

17. Mueller SG, Laxer KD, Scanlon C, et al. Different structural correlates for verbal memory impairment in temporal lobe epilepsy with and without mesial temporal lobe sclerosis. Human brain mapping. 2012;33(2):489–499.

18. Oyegbile T, Hansen R, Magnotta V, et al. Quantitative measurement of cortical surface features in localization-related temporal lobe epilepsy. Neuropsychology. 2004;18(4):729.

19. Dabbs K, Jones J, Seidenberg M, Hermann B. Neuroanatomical correlates of cognitive phenotypes in temporal lobe epilepsy. Epilepsy & Behavior. 2009/08/01/ 2009;15(4):445–451. doi:https://doi.org/10.1016/j.yebeh.2009.05.012

20. Rodríguez-Cruces R, Bernhardt BC, Concha L. Multidimensional associations between cognition and connectome organization in temporal lobe epilepsy. NeuroImage. 2020;213:116706.

21. Reyes A, Kaestner E, Bahrami N, et al. Cognitive phenotypes in temporal lobe epilepsy are associated with distinct patterns of white matter network abnormalities. Neurology. 2019;92(17):e1957–e1968.

22. Hermann B, Conant LL, Cook CJ, et al. Network, clinical and sociodemographic features of cognitive phenotypes in temporal lobe epilepsy. NeuroImage: Clinical. 2020/01/01/ 2020;27:102341. doi:https://doi.org/10.1016/j.nicl.2020.102341

23. Fadaie F, Lee HM, Caldairou B, et al. Atypical functional connectome hierarchy impacts cognition in temporal lobe epilepsy. Epilepsia. 2021;62(11):2589–2603.

24. Girardi-Schappo M, Fadaie F, Lee HM, et al. Altered communication dynamics reflect cognitive deficits in temporal lobe epilepsy. Epilepsia. 2021;62(4):1022–1033. doi:https://doi.org/10.1111/epi.16864

25. Tailby C, Kowalczyk MA, Jackson GD. Cognitive impairment in epilepsy: the role of reduced network flexibility. Annals of Clinical and Translational Neurology. 2018;5(1):29–40. doi:https://doi.org/10.1002/acn3.503

26. Margulies DS, Ghosh SS, Goulas A, et al. Situating the default-mode network along a principal gradient of macroscale cortical organization. Proceedings of the National Academy of Sciences. 2016;113(44):12574–12579.

27. Smallwood J, Bernhardt BC, Leech R, Bzdok D, Jefferies E, Margulies DS. The default mode network in cognition: a topographical perspective. Nature reviews neuroscience. 2021;22(8):503–513.

28. Moscovitch M, Cabeza R, Winocur G, Nadel L. Episodic memory and beyond: the hippocampus and neocortex in transformation. Annual review of psychology. 2016;67:105.

29. Paquola C, Bethlehem RA, Seidlitz J, et al. Shifts in myeloarchitecture characterise adolescent development of cortical gradients. Elife. 2019;8:e50482.

30. Paquola C, Seidlitz J, Benkarim O, et al. A multi-scale cortical wiring space links cellular architecture and functional dynamics in the human brain. PLoS biology. 2020;18(11):e3000979.

31. Royer J, Paquola C, Larivière S, et al. Myeloarchitecture gradients in the human insula: Histological underpinnings and association to intrinsic functional connectivity. Neuroimage. 2020;216:116859.

32. Paquola C, Vos De Wael R, Wagstyl K, et al. Microstructural and functional gradients are increasingly dissociated in transmodal cortices. PLoS biology. 2019;17(5):e3000284.

33. Stüber C, Morawski M, Schäfer A, et al. Myelin and iron concentration in the human brain: a quantitative study of MRI contrast. Neuroimage. 2014;93:95–106.

34. Mesulam M-M. From sensation to cognition. Brain: a journal of neurology. 1998;121(6):1013–1052.

35. Valk SL, Xu T, Paquola C, et al. Genetic and phylogenetic uncoupling of structure and function in human transmodal cortex. bioRxiv. 2021:2021.06.08.447522. doi:10.1101/2021.06.08.447522

36. Murphy C, Jefferies E, Rueschemeyer S-A, et al. Distant from input: Evidence of regions within the default mode network supporting perceptually-decoupled and conceptually-guided cognition. NeuroImage. 2018/05/01/ 2018;171:393–401. doi:https://doi.org/10.1016/j.neuroimage.2018.01.017

37. Murphy C, Wang H-T, Konu D, et al. Modes of operation: A topographic neural gradient supporting stimulus dependent and independent cognition. NeuroImage. 2019/02/01/ 2019;186:487–496. doi:https://doi.org/10.1016/j.neuroimage.2018.11.009

38. Zhang M, Bernhardt BC, Wang X, et al. Perceptual coupling and decoupling of the default mode network during mind-wandering and reading. Elife. 2022;11:e74011.

39. Zhang M, McNab F, Smallwood J, Jefferies E. Perceptual coupling and decoupling are associated with individual differences in working memory encoding and maintenance. Cerebral Cortex. 2022;32(18):3959–3974.

40. Engel Jr J. Outcome with respect to epileptic seizures. Surgical treatment of the epilepsies. 1993:609–621.

41. Blümcke I, Thom M, Aronica E, et al. International consensus classification of hippocampal sclerosis in temporal lobe epilepsy: a Task Force report from the ILAE Commission on Diagnostic Methods. Epilepsia. 2013;54(7):1315–1329.

42. Royer J, Rodríguez-Cruces R, Tavakol S, et al. An open MRI dataset for multiscale neuroscience. bioRxiv. 2021;

43. Haast RA, Ivanov D, Formisano E, Uludağ K. Reproducibility and reliability of quantitative and weighted T1 and T2* mapping for myelin-based cortical parcellation at 7 Tesla. Frontiers in neuroanatomy. 2016;10:112.

44. Marques JP, Kober T, Krueger G, van der Zwaag W, Van de Moortele P-F, Gruetter R. MP2RAGE, a self bias-field corrected sequence for improved segmentation and T1-mapping at high field. Neuroimage. 2010;49(2):1271–1281.

45. Cruces RR, Royer J, Herholz P, et al. Micapipe: A Pipeline for Multimodal Neuroimaging and Connectome Analysis. bioRxiv. 2022:2022.01.31.478189. doi:10.1101/2022.01.31.478189

46. Dale AM, Fischl B, Sereno MI. Cortical surface-based analysis: I. Segmentation and surface reconstruction. Neuroimage. 1999;9(2):179–194.

47. Fischl B, Sereno MI, Dale AM. Cortical surface-based analysis: II: inflation, flattening, and a surface-based coordinate system. Neuroimage. 1999;9(2):195–207.

48. Fischl B, Sereno MI, Tootell RB, Dale AM. High-resolution intersubject averaging and a coordinate system for the cortical surface. Human brain mapping. 1999;8(4):272–284.

49. Cox RW. AFNI: software for analysis and visualization of functional magnetic resonance neuroimages. Computers and Biomedical research. 1996;29(3):162–173.

50. Jenkinson M, Beckmann CF, Behrens TE, Woolrich MW, Smith SM. Fsl. Neuroimage. 2012;62(2):782–790.

51. Salimi-Khorshidi G, Douaud G, Beckmann CF, Glasser MF, Griffanti L, Smith SM. Automatic denoising of functional MRI data: combining independent component analysis and hierarchical fusion of classifiers. Neuroimage. 2014;90:449–468.

52. Greve DN, Fischl B. Accurate and robust brain image alignment using boundary-based registration. Neuroimage. 2009;48(1):63–72.

53. Marcus DS, Harms MP, Snyder AZ, et al. Human Connectome Project informatics: Quality control, database services, and data visualization. NeuroImage. 2013/10/15/ 2013;80:202–219. doi:https://doi.org/10.1016/j.neuroimage.2013.05.077

54. Glasser MF, Sotiropoulos SN, Wilson JA, et al. The minimal preprocessing pipelines for the Human Connectome Project. NeuroImage. 2013/10/15/ 2013;80:105–124. doi:https://doi.org/10.1016/j.neuroimage.2013.04.127

55. Waehnert MD, Dinse J, Schäfer A, et al. A subject-specific framework for in vivo myeloarchitectonic analysis using high resolution quantitative MRI. Neuroimage. 2016;125:94–107.

56. Coifman RR, Lafon S, Lee AB, et al. Geometric diffusions as a tool for harmonic analysis and structure definition of data: Diffusion maps. Proceedings of the national academy of sciences. 2005;102(21):7426–7431.

57. Hong S-J, De Wael RV, Bethlehem RA, et al. Atypical functional connectome hierarchy in autism. Nature communications. 2019;10(1):1–13.

58. Larivière S, Vos de Wael R, Hong S-J, et al. Multiscale structure–function gradients in the neonatal connectome. Cerebral Cortex. 2020;30(1):47–58.

59. Vos de Wael R, Benkarim O, Paquola C, et al. BrainSpace: a toolbox for the analysis of macroscale gradients in neuroimaging and connectomics datasets. Communications biology. 2020;3(1):1–10.

60. Liu M, Bernhardt BC, Hong S-J, Caldairou B, Bernasconi A, Bernasconi N. The superficial white matter in temporal lobe epilepsy: a key link between structural and functional network disruptions. Brain. 2016;139(9):2431–2440.

61. Worsley KJ, Taylor J, Carbonell F, et al. A Matlab toolbox for the statistical analysis of univariate and multivariate surface and volumetric data using linear mixed effects models and random field theory. 2009:S102.

62. Scholtens LH, de Reus MA, de Lange SC, Schmidt R, van den Heuvel MP. An mri von economo–koskinas atlas. NeuroImage. 2018;170:249–256.

63. von Economo CF, Koskinas GN. Die cytoarchitektonik der hirnrinde des erwachsenen menschen. J. Springer; 1925.

64. Larivière S, Paquola C, Park B-y, et al. The ENIGMA Toolbox: multiscale neural contextualization of multisite neuroimaging datasets. Nature Methods. 2021;18(7):698–700.

65. Mesulam M-M. Principles of behavioral and cognitive neurology. Oxford University Press; 2000.

66. Amunts K, Lepage C, Borgeat L, et al. BigBrain: an ultrahigh-resolution 3D human brain model. Science. 2013;340(6139):1472–1475.

67. Paquola C, Royer J, Lewis LB, et al. The BigBrainWarp toolbox for integration of BigBrain 3D histology with multimodal neuroimaging. eLife. 2021;10:e70119.

68. Fornito A, Arnatkevičiūtė A, Fulcher BD. Bridging the gap between connectome and transcriptome. Trends in Cognitive Sciences. 2019;23(1):34–50.

69. Hawrylycz MJ, Lein ES, Guillozet-Bongaarts AL, et al. An anatomically comprehensive atlas of the adult human brain transcriptome. Nature. 2012;489(7416):391–399.

70. Larivière S, Royer J, Rodríguez-Cruces R, et al. Structural network alterations in focal and generalized epilepsy assessed in a worldwide ENIGMA study follow axes of epilepsy risk gene expression. Nature Communications. 2022/07/27 2022;13(1):4320. doi:10.1038/s41467-022-31730-5

71. Schaefer A, Kong R, Gordon EM, et al. Local-global parcellation of the human cerebral cortex from intrinsic functional connectivity MRI. Cerebral cortex. 2018;28(9):3095–3114.

72. Consortium TILAE. Genome-wide mega-analysis identifies 16 loci and highlights diverse biological mechanisms in the common epilepsies. Nature communications. 2018;9

73. Larivière S, Rodríguez-Cruces R, Royer J, et al. Network-based atrophy modeling in the common epilepsies: A worldwide ENIGMA study. Science advances. 2020;6(47):eabc6457.

74. Whelan CD, Altmann A, Botía JA, et al. Structural brain abnormalities in the common epilepsies assessed in a worldwide ENIGMA study. Brain. 2018;141(2):391–408.

75. Garbelli R, Milesi G, Medici V, et al. Blurring in patients with temporal lobe epilepsy: clinical, high-field imaging and ultrastructural study. Brain. 2012;135(8):2337–2349.

76. Meyer A, Falconer MA, Beck E. Pathological findings in temporal lobe epilepsy. Journal of neurology, neurosurgery, and psychiatry. 1954;17(4):276.

77. Cavanagh J, Meyer A. Aetiological aspects of Ammon’s horn sclerosis associated with temporal lobe epilepsy. British medical journal. 1956;2(5006):1403.

78. Bruton CJ. The neuropathology of temporal lobe epilepsy. Maudsley monographs. 1988;31

79. Blanc F, Martinian L, Liagkouras I, Catarino C, Sisodiya SM, Thom M. Investigation of widespread neocortical pathology associated with hippocampal sclerosis in epilepsy: a postmortem study. Epilepsia. 2011;52(1):10–21.

80. Bernhardt BC, Fadaie F, de Wael RV, et al. Preferential susceptibility of limbic cortices to microstructural damage in temporal lobe epilepsy: A quantitative T1 mapping study. Neuroimage. 2018;182:294–303.

81. Winston GP, Vos SB, Caldairou B, et al. Microstructural imaging in temporal lobe epilepsy: Diffusion imaging changes relate to reduced neurite density. NeuroImage: Clinical. 2020;26:102231.

82. Thom M, Holton J, D’Arrigo C, et al. Microdysgenesis with abnormal cortical myelinated fibres in temporal lobe epilepsy: a histopathological study with calbindin D-28-K immunohistochemistry. Neuropathology and applied neurobiology. 2000;26(3):251–257.

83. Hardiman O, Burke T, Phillips J, et al. Microdysgenesis in resected temporal neocortex: incidence and clinical significance in focal epilepsy. Neurology. 1988;38(7):1041–1041.

84. Eriksson SH, Nordborg C, Thom M, Sisodiya SM. Microdysgenesis in mesial temporal lobe epilepsy. Annals of neurology. 2004;55(4):596–597.

85. Kasper BS, Stefan H, Paulus W. Microdysgenesis in mesial temporal lobe epilepsy: a clinicopathological study. Annals of Neurology: Official Journal of the American Neurological Association and the Child Neurology Society. 2003;54(4):501–506.

86. Rakic P. Neurogenesis in adult primate neocortex: an evaluation of the evidence. Nature Reviews Neuroscience. 2002/01/01 2002;3(1):65–71. doi:10.1038/nrn700

87. Haas CA, Dudeck O, Kirsch M, et al. Role for reelin in the development of granule cell dispersion in temporal lobe epilepsy. Journal of Neuroscience. 2002;22(14):5797–5802.

88. Machado RA, Benjumea-Cuartas V, Zapata Berruecos JF, Agudelo-Flóres PM, Salazar-Peláez LM. Reelin, tau phosphorylation and psychiatric complications in patients with hippocampal sclerosis and structural abnormalities in temporal lobe epilepsy. Epilepsy & Behavior. 2019/07/01/ 2019;96:192–199. doi:https://doi.org/10.1016/j.yebeh.2019.04.052

89. Siebzehnrubl FA, Blumcke I. Neurogenesis in the human hippocampus and its relevance to temporal lobe epilepsies. Epilepsia. 2008;49:55–65.

90. Heinrich C, Nitta N, Flubacher A, et al. Reelin deficiency and displacement of mature neurons, but not neurogenesis, underlie the formation of granule cell dispersion in the epileptic hippocampus. Journal of Neuroscience. 2006;26(17):4701–4713.

91. García-Cabezas MÁ, Zikopoulos B, Barbas H. The Structural Model: a theory linking connections, plasticity, pathology, development and evolution of the cerebral cortex. Brain Structure and Function. 2019;224(3):985–1008.

92. Barbas H. General cortical and special prefrontal connections: principles from structure to function. Annual review of neuroscience. 2015;38:269–289.

93. Goulas A, Zilles K, Hilgetag CC. Cortical gradients and laminar projections in mammals. Trends in Neurosciences. 2018;41(11):775–788.

94. Larivière S, Weng Y, Vos de Wael R, et al. Functional connectome contractions in temporal lobe epilepsy: Microstructural underpinnings and predictors of surgical outcome. Epilepsia. 2020;61(6):1221–1233.

95. Coito A, Genetti M, Pittau F, et al. Altered directed functional connectivity in temporal lobe epilepsy in the absence of interictal spikes: a high density EEG study. Epilepsia. 2016;57(3):402–411.

96. Royer J, Bernhardt BC, Larivière S, et al. Epilepsy and brain network hubs. Epilepsia. 2022;63(3):537–550. doi:https://doi.org/10.1111/epi.17171

97. Bernhardt BC, Fadaie F, Liu M, et al. Temporal lobe epilepsy: Hippocampal pathology modulates connectome topology and controllability. Neurology. 2019;92(19):e2209–e2220.

98. Gleichgerrcht E, Keller SS, Drane DL, et al. Temporal lobe epilepsy surgical outcomes can be inferred based on structural connectome hubs: a machine learning study. Annals of neurology. 2020;88(5):970–983.

99. Betzel RF, Bassett DS. Multi-scale brain networks. Neuroimage. 2017;160:73–83.

100. Huntenburg JM, Bazin P-L, Margulies DS. Large-scale gradients in human cortical organization. Trends in cognitive sciences. 2018;22(1):21–31.

101. Fulcher BD, Murray JD, Zerbi V, Wang X-J. Multimodal gradients across mouse cortex. Proceedings of the National Academy of Sciences. 2019;116(10):4689–4695.

102. Pandya D, Petrides M, Cipolloni PB. Cerebral cortex: architecture, connections, and the dual origin concept. Oxford University Press; 2015.

103. García-Cabezas MÁ, Joyce MKP, John YJ, Zikopoulos B, Barbas H. Mirror trends of plasticity and stability indicators in primate prefrontal cortex. European Journal of Neuroscience. 2017;46(8):2392–2405.

104. Neve RL, Finch EA, Bird ED, Benowitz LI. Growth-associated protein GAP-43 is expressed selectively in associative regions of the adult human brain. Proceedings of the National Academy of Sciences. 1988;85(10):3638–3642.

105. Benowitz LI, Routtenberg A. GAP-43: an intrinsic determinant of neuronal development and plasticity. Trends in neurosciences. 1997;20(2):84–91.

106. Kinjo ER, Rodríguez PXR, Dos Santos BA, et al. New insights on temporal lobe epilepsy based on plasticity-related network changes and high-order statistics. Molecular Neurobiology. 2018;55(5):3990–3998.

107. Tolner EA, Van Vliet EA, Holtmaat AJ, et al. GAP-43 mRNA and protein expression in the hippocampal and parahippocampal region during the course of epileptogenesis in rats. European Journal of Neuroscience. 2003;17(11):2369–2380.

108. Elmér E, Kokaia M, Kokaia Z, Ferencz I, Lindvall O. Delayed kindling development after rapidly recurring seizures: relation to mossy fiber sprouting and neurotrophin, GAP-43 and dynorphin gene expression. Brain research. 1996;712(1):19–34.

109. Aigner L, Arber S, Kapfhammer JP, et al. Overexpression of the neural growth-associated protein GAP-43 induces nerve sprouting in the adult nervous system of transgenic mice. Cell. 1995;83(2):269–278.

110. Schleicher A, Amunts K, Geyer S, Morosan P, Zilles K. Observer-independent method for microstructural parcellation of cerebral cortex: a quantitative approach to cytoarchitectonics. Neuroimage. 1999;9(1):165–177.

111. Schleicher A, Palomero-Gallagher N, Morosan P, et al. Quantitative architectural analysis: a new approach to cortical mapping. Anatomy and embryology. 2005;210(5):373–386.

112. Goubran M, Bernhardt BC, Cantor-Rivera D, et al. In vivo MRI signatures of hippocampal subfield pathology in intractable epilepsy. Human brain mapping. 2016;37(3):1103–1119.

113. Goubran M, Hammond RR, de Ribaupierre S, et al. Magnetic resonance imaging and histology correlation in the neocortex in temporal lobe epilepsy. Annals of neurology. 2015;77(2):237–250.

114. Burt JB, Demirtaş M, Eckner WJ, et al. Hierarchy of transcriptomic specialization across human cortex captured by structural neuroimaging topography. Nature neuroscience. 2018;21(9):1251–1259.

115. Darmanis S, Sloan SA, Zhang Y, et al. A survey of human brain transcriptome diversity at the single cell level. Proceedings of the National Academy of Sciences. 2015;112(23):7285–7290.

116. He Z, Han D, Efimova O, et al. Comprehensive transcriptome analysis of neocortical layers in humans, chimpanzees and macaques. Nature neuroscience. 2017;20(6):886–895.

117. Bernard A, Lubbers LS, Tanis KQ, et al. Transcriptional architecture of the primate neocortex. Neuron. 2012;73(6):1083–1099.

118. Belgard TG, Marques AC, Oliver PL, et al. A transcriptomic atlas of mouse neocortical layers. Neuron. 2011;71(4):605–616.

119. Krienen FM, Yeo BT, Ge T, Buckner RL, Sherwood CC. Transcriptional profiles of supragranular-enriched genes associate with corticocortical network architecture in the human brain. Proceedings of the National Academy of Sciences. 2016;113(4):E469–E478.

120. Wray NR, Yang J, Hayes BJ, Price AL, Goddard ME, Visscher PM. Pitfalls of predicting complex traits from SNPs. Nature Reviews Genetics. 2013;14(7):507–515.

121. Rodríguez-Cruces R, Velázquez-Pérez L, Rodríguez-Leyva I, et al. Association of white matter diffusion characteristics and cognitive deficits in temporal lobe epilepsy. Epilepsy & Behavior. 2018;79:138–145.

122. Fletcher PC, Henson RNA. Frontal lobes and human memory: insights from functional neuroimaging. Brain. 2001;124(5):849–881.

123. Simons JS, Spiers HJ. Prefrontal and medial temporal lobe interactions in long-term memory. Nature reviews neuroscience. 2003;4(8):637–648.

124. Sidhu MK, Stretton J, Winston GP, et al. A functional magnetic resonance imaging study mapping the episodic memory encoding network in temporal lobe epilepsy. Brain. 2013;136(6):1868–1888. doi:10.1093/brain/awt099

125. Alessio A, Pereira FR, Sercheli MS, et al. Brain plasticity for verbal and visual memories in patients with mesial temporal lobe epilepsy and hippocampal sclerosis: an fMRI study. Human brain mapping. 2013;34(1):186–199.

126. Maccotta L, Buckner RL, Gilliam FG, Ojemann JG. Changing frontal contributions to memory before and after medial temporal lobectomy. Cerebral Cortex. 2007;17(2):443–456.

127. Richardson MP, Strange BA, Duncan JS, Dolan RJ. Preserved verbal memory function in left medial temporal pathology involves reorganisation of function to right medial temporal lobe. Neuroimage. 2003;20:S112–S119.

128. Fleury M, Buck S, Binding LP, et al. Episodic memory network connectivity in temporal lobe epilepsy. Epilepsia. 2022;

129. Dupont S, Van de Moortele PF, Samson S, et al. Episodic memory in left temporal lobe epilepsy: a functional MRI study. Brain. 2000;123(8):1722–1732. doi:10.1093/brain/123.8.1722

130. Ives-Deliperi V, Butler JT. Mechanisms of cognitive impairment in temporal lobe epilepsy: a systematic review of resting-state functional connectivity studies. Epilepsy & Behavior. 2021;115:107686.

