## Supplementary materials for "Cortical microstructural gradients capture memory network reorganization in temporal lobe epilepsy"

### SUPPLEMENTARY MATERIAL

### CORRESPONDING AUTHORS

Jessica Royer, Psy.D.


Boris C. Bernhardt, PhD


### SUPPLEMENTARY METHODS AND RESULTS

*Quantitative intracortical intensity profiling*

We investigated the laminar underpinnings of microstructural gradient reconfigurations using quantitative profiling of intracortical R1 intensities across cortical depths. This approach characterizes the shape of intracortical microstructural profiles by calculating the central moments (mean amplitude, mean depth of peak, SD, skewness, and kurtosis) of intensity values distributed between the pial and white matter boundaries (**FIGURE S4A**). These features thus captured subject-specific properties of local laminar myeloarchitecture across the cortex. Indeed, these metrics could differentiate primary and unimodal cortices from heteromodal and paralimbic regions (**FIGURE S4B**). Each profile shape map was statistically compared between TLE and HCs using surface-based linear models, while controlling for effects of age and sex (**FIGURE S5A**). We then computed spatial correlations between effect size maps associated with each statistical moment and gradient contractions (**FIGURE S5B**). Statistical significance was determined using nonparametric spin permutation testing (1,000 permutations) implemented in the BrainSpace toolbox. Targeted post-hoc analyses then compared average microstructural moment changes within significant clusters of gradient contractions in TLE patients relative to controls (**FIGURE S5C**). Compared to controls, changes in each central moment could be observed in the TLE group: Although these alterations were not confined to ipsilateral temporal cortices, consistent alterations in all moments could be observed in this region.

*Controlling for cortical morphology and pericortical blurring*

We repeated group comparisons in microstructural gradients while additionally controlling for cortical thickness. For this purpose, we fit a vertex-wise linear model predicting individual microstructural gradients from cortical thickness, with residuals serving as estimates of thickness-corrected microstructural gradients. [1]. Controlling for cortical thickness at the vertex-level had little impact on group differences in gradient scores (**FIGURE S6A**). In line with initial findings, correcting individual gradient maps for vertex-wise measures of cortical thickness revealed significant microstructural gradient contractions in ipsilateral temporopolar and lateral prefrontal regions ($p_{\mathrm{FWE}}<0.001$). We additionally controlled for changed in grey-white matter interface blurring. We estimated blurring from qT1 intensity gradients computed perpendicular to the cortical mantle, in line with previous work [1-3]. The same vertex-level procedure as for cortical thickness was applied to correct for cortex-wide measurements of cortical blurring. Results were also consistent with initial findings after correcting for vertex-wise grey-white matter interface contrast measured from qT1 images (**FIGURE S6B**).

**TABLE S1**

| ID | Age | Sex | HD | TLE Laterality | MRI diagnosis | Surgery | Engel outcome |
|---|---|---|---|---|---|---|---|
| PX002 | 24 | M | R | R | T2/FLAIR hyperintensity in hippocampus | NA | NA |
| PX003 | 54 | M | R | L | T2/FLAIR hyperintensity in hippocampus | NA | NA |
| PX005 | 26 | F | R | L | MRI-negative | RFTC | 1a |
| PX008 | 20 | F | R | L | Tumour in the uncus | Tumour resection | 1a |
| PX012 | 33 | F | R | L | MTS | NA | NA |
| PX013 | 57 | M | R | L | MTS | SAH | |
| PX019 | 31 | M | R | L | Suspected FCD in parahippocampal gyrus | NA | NA |
| PX021 | 55 | F | R | L | T2/FLAIR hyperintensity in hippocampus | SAH | 1a |
| PX025 | 25 | M | R | R | T2/FLAIR hyperintensity in hippocampus | NA | NA |
| PX027 | 40 | F | L | L | MRI-negative | NA | NA |
| PX028 | 22 | F | R | R | MRI-negative | RFTC; CAH | 1a |
| PX032 | 53 | F | R | L | MTS | NA | NA |
| PX036 | 24 | M | R | L | Cystic malformation and FCD | NA | NA |
| PX042 | 38 | M | R | L | MRI-negative | CAH | 1a |
| PX043 | 37 | F | R | R | MTS | NA | NA |
| PX044 | 43 | M | R | R | MTS | NA | NA |
| PX047 | 35 | M | R | L | MRI negative | CAH | 1a |
| PX050 | 32 | M | R | R | MRI negative | RFTC | 1a |
| PX053 | 41 | M | R | L | T2/FLAIR hyperintensity in mesio-temporal lobe and insula | RFTC | 4 |
| PX055 | 27 | F | R | L | MRI-negative | NA | NA |
| PX062 | 44 | M | R | L | MTS | CAH | 1a |

**TABLE S1.** Detailed clinical information of discovery cohort. Abbreviations: *CAH*: cortico-amygdalohippocampectomy; *FCD*: Focal cortical dysplasia; *HD*: Handedness; *MTS*: mesio-temporal sclerosis; *RFTC*: stereotactic radiofrequency thermocoagulation; *SAH*: selective amygdalo-hippocampectomy

**TABLE S2**

| Disorder | Genes |
|---|---|
| All epilepsies | BCL11A, BRD7, FANCL, HEATR3, SCN1A, SCN2A, SCN3A, TTC21B |
| Childhood absence epilepsy | BCL11A, FANCL, ZEB2 |
| Focal epilepsy | SCN1A, SCN2A, SCN3A, TTC21B |
| Focal epilepsy (hippocampal sclerosis) | C3orf33, GJA1, KCNAB1, SLC33A1 |
| Generalized epilepsy | ATXN1, BCL11A, FANCL, GABRA2, GRIK1, KCNN2, PCDH7, PNPO, SCN1A, SCN2A, SCN3A, STAT4, TTC21B |
| Juvenile myoclonic epilepsy | STX1B |

**TABLE S2.** List of disease-specific risk genes used in the present study extracted from previously published GWAS [4]. Gene lists and pre-processed gene expression data [5] are openly available as part of the ENIGMA toolbox [6].

**FIGURE S1**

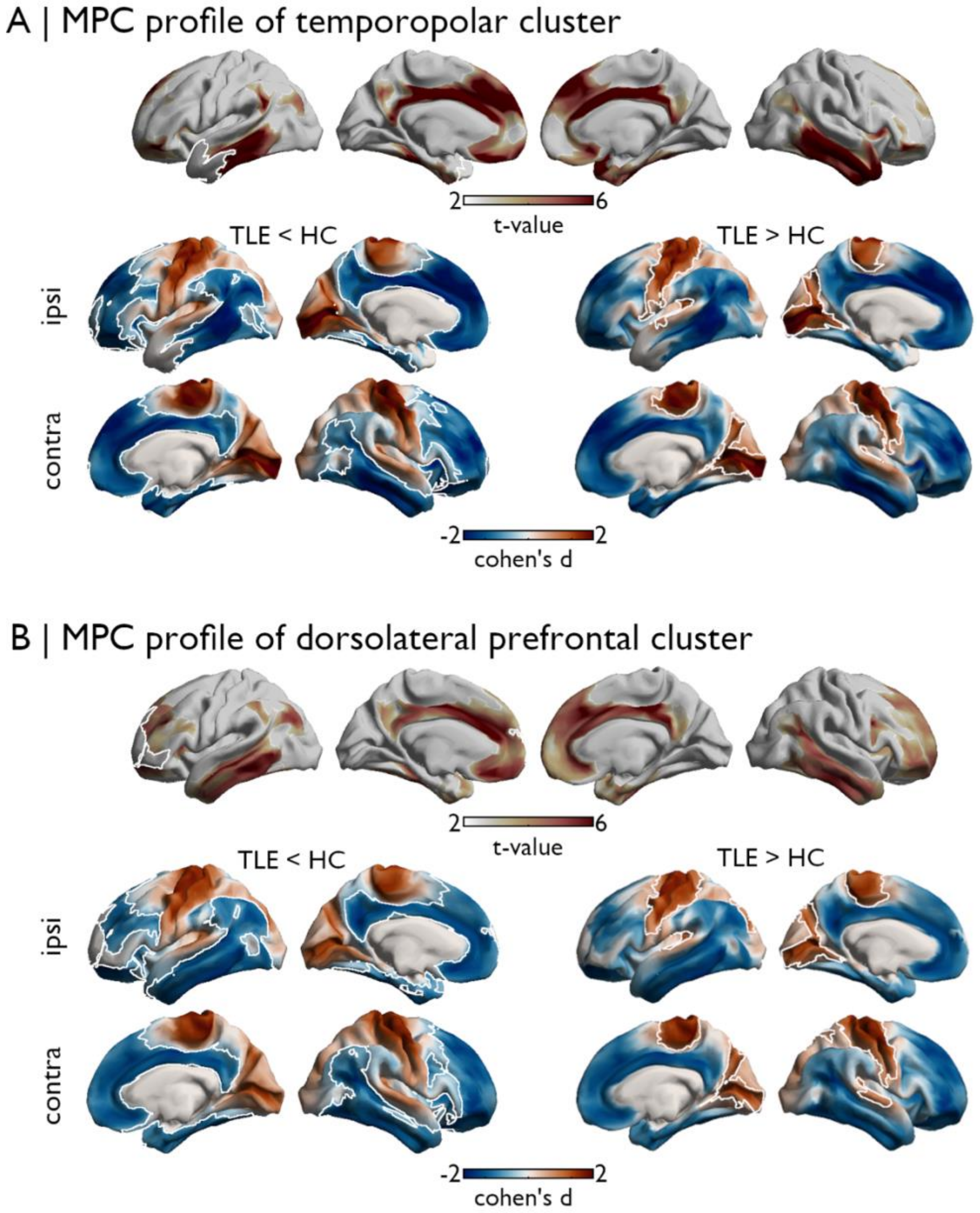


**Figure S1. *Post hoc* analysis of microstructural profile similarity patterns. (A)** The top panel shows the microstructural similarity profile of vertices located in the temporopolar cluster for the HC group (average of vertices in the cluster, which is outlined in white). Compared to controls, the TLE group showed reduced microstructural similarity to paralimbic and transmodal cortices (left, significant regions outlined in white, $p$FWE < 0.025), and higher microstructural profile similarity to unimodal sensory and motor regions (right, significant regions outlined in white, $p$FWE < 0.025). **(B)** The top panel shows the microstructural similarity profile of vertices located in the prefrontal cluster for the HC group (average of vertices in the cluster, which is outlined in white). Compared to controls, the TLE group showed reduced microstructural similarity to paralimbic and transmodal cortices (left, significant regions outlined in white, $p$FWE < 0.025), and higher microstructural profile similarity to unimodal sensory and motor regions (right, significant regions outlined in white, $p$FWE < 0.025).

**FIGURE S2**

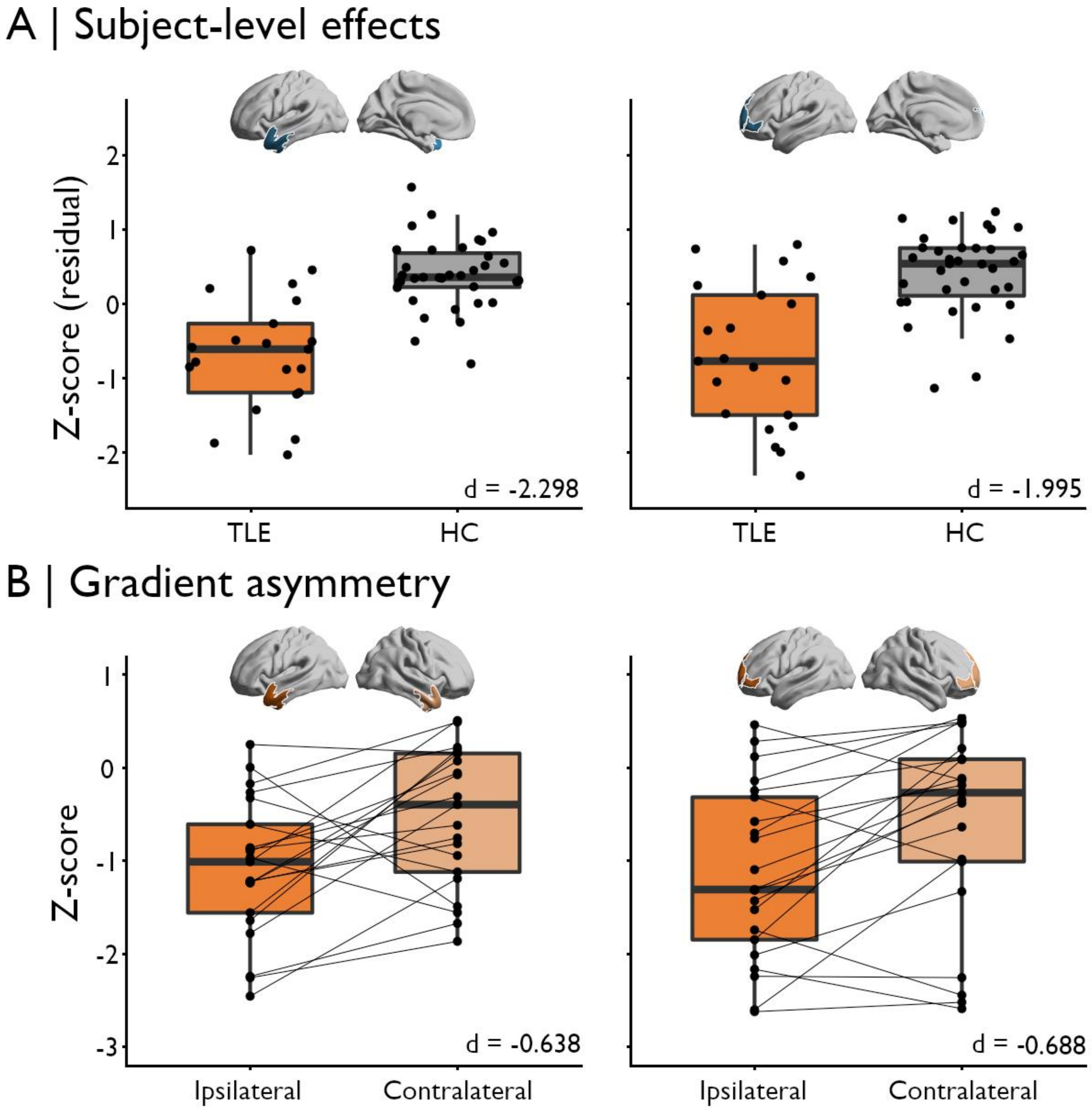


**Figure S2. Patient-level consistency of microstructural gradient contractions and asymmetry. (A)** 80.95% of patients showed moderately negative average z-scores ($z$<-1) in either significant cluster of gradient reductions (57.14% at $z$<-1.5, 33.33% at $z$<-2). After correction for age and sex, we observed strong reductions in gradient scores in TLE patients compared to HCs in both temporal lobe (cohen's $d$=-2.298) and prefrontal clusters (cohen's $d$=-1.995). **(B)** Microstructural gradient contractions also showed potential for lateralization of the seizure focus in TLE. Indeed, 90.48% of patients showed lower average z-scores in the ipsilateral vs. contralateral hemisphere in either significant cluster of gradient reductions. When considering each cluster independently, this proportion remained high, with 76.14% of patients showing lower average z-scores in the ipsilateral temporal (cohen's $d$=-0.638) or prefrontal cluster (cohen's $d$=-0.688) relative to its homologous region in the contralateral hemisphere.

### FIGURE S3

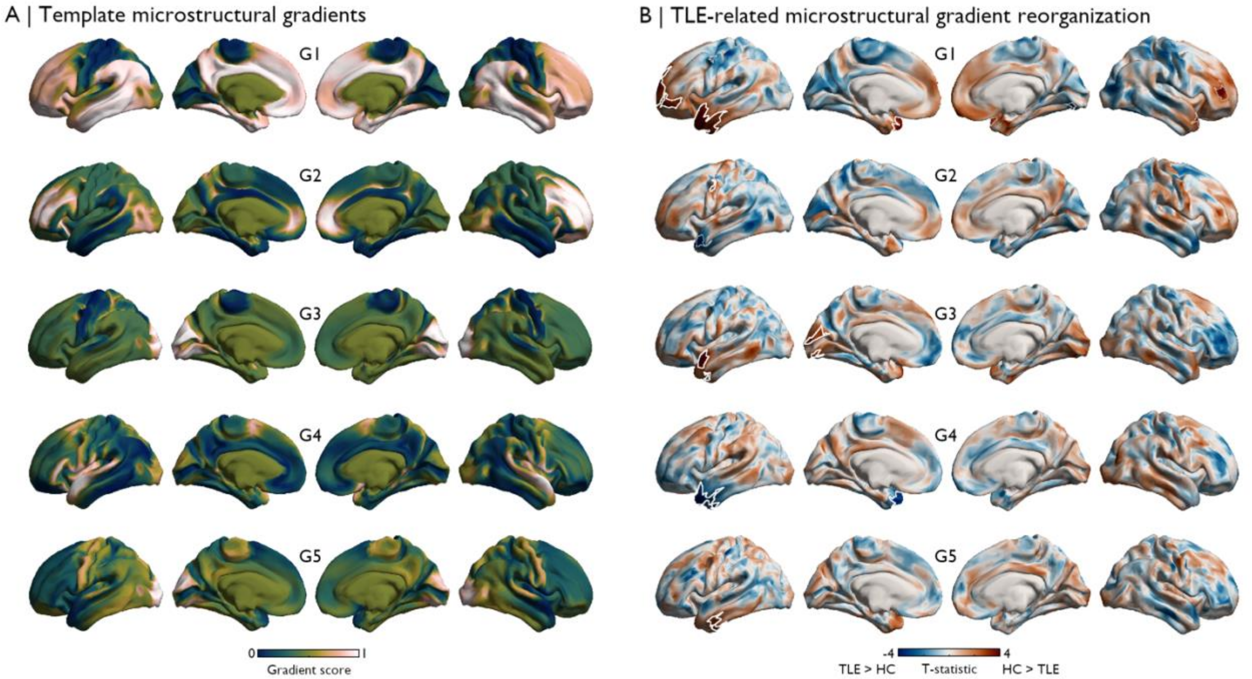


**Figure S3. Topography of TLE and HC microstructural gradients 1-5. (A)** Sample-wide (including all TLE and HC participants) gradient templates (G1-G5). **(B)** Differences in gradient scores between TLE and HCs are shown in unthresholded t-statistic maps, with significant clusters of findings outlined in white ($p_{FWE}<0.025$) and trend-level effects outlined in thinner grey clusters ($p_{FWE}<0.1$). As in our main analyses, all group comparisons corrected for age and sex. Anterior temporal regions ipsilateral to the seizure focus were systematically perturbed in TLE relative to corresponding gradients in HCs.

**FIGURE S4**

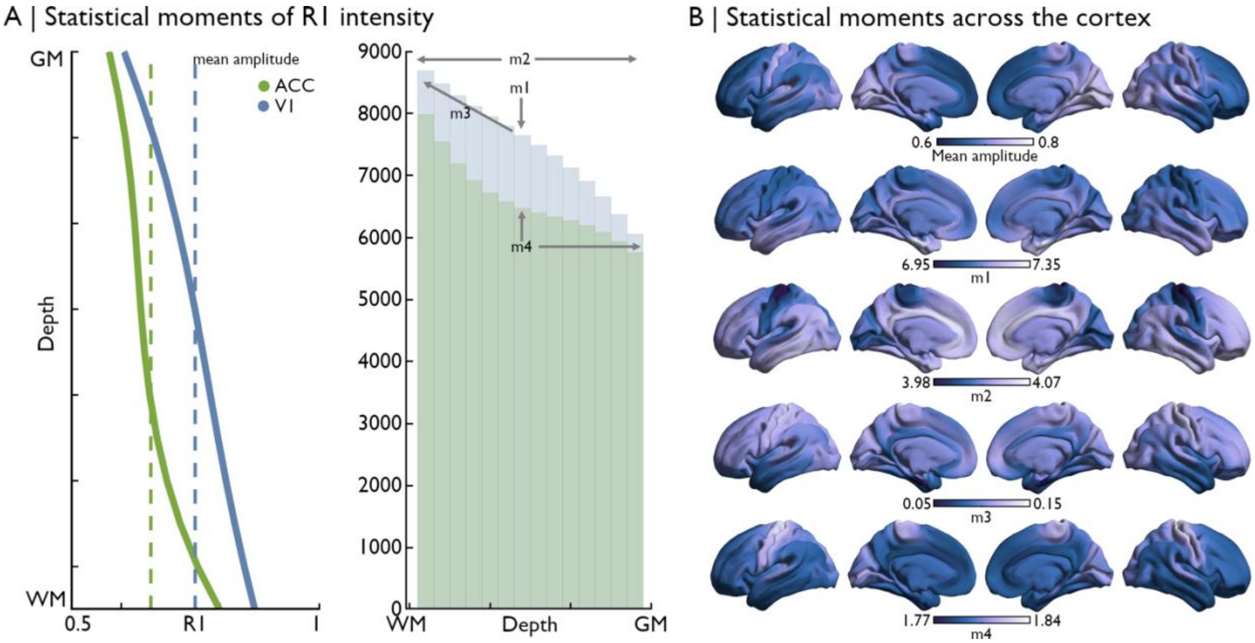


**Figure S4. Group-level central moment maps. (A)** Intensity profiles sampled at a single vertex in the anterior cingulate cortex (ACC, green) and primary visual cortex (V1, blue), averaged across all healthy control participants. The mean amplitude of each profile is indicated by the dashed line, and was computed by averaging the intensity values sampled at each depth for each vertex. The shape of each profile could be quantified by first converting each profile to a frequency distribution: The corresponding frequency distribution of the ACC and V1 profiles (left) are displayed in a histogram (right). We computed the mean (m1), standard deviation (m2), skewness (m3), and kurtosis (m4) of depth-dependent R1 intensity distributions, thus capturing the overall shape of vertex-wise microstructural profiles along the cortical sheet. **(B)** Central moment maps were averaged across all subjects and projected to the cortical surface. In line with previous work, these metrics could differentiate primary and unimodal cortices from heteromodal and paralimbic regions.

**FIGURE S5**

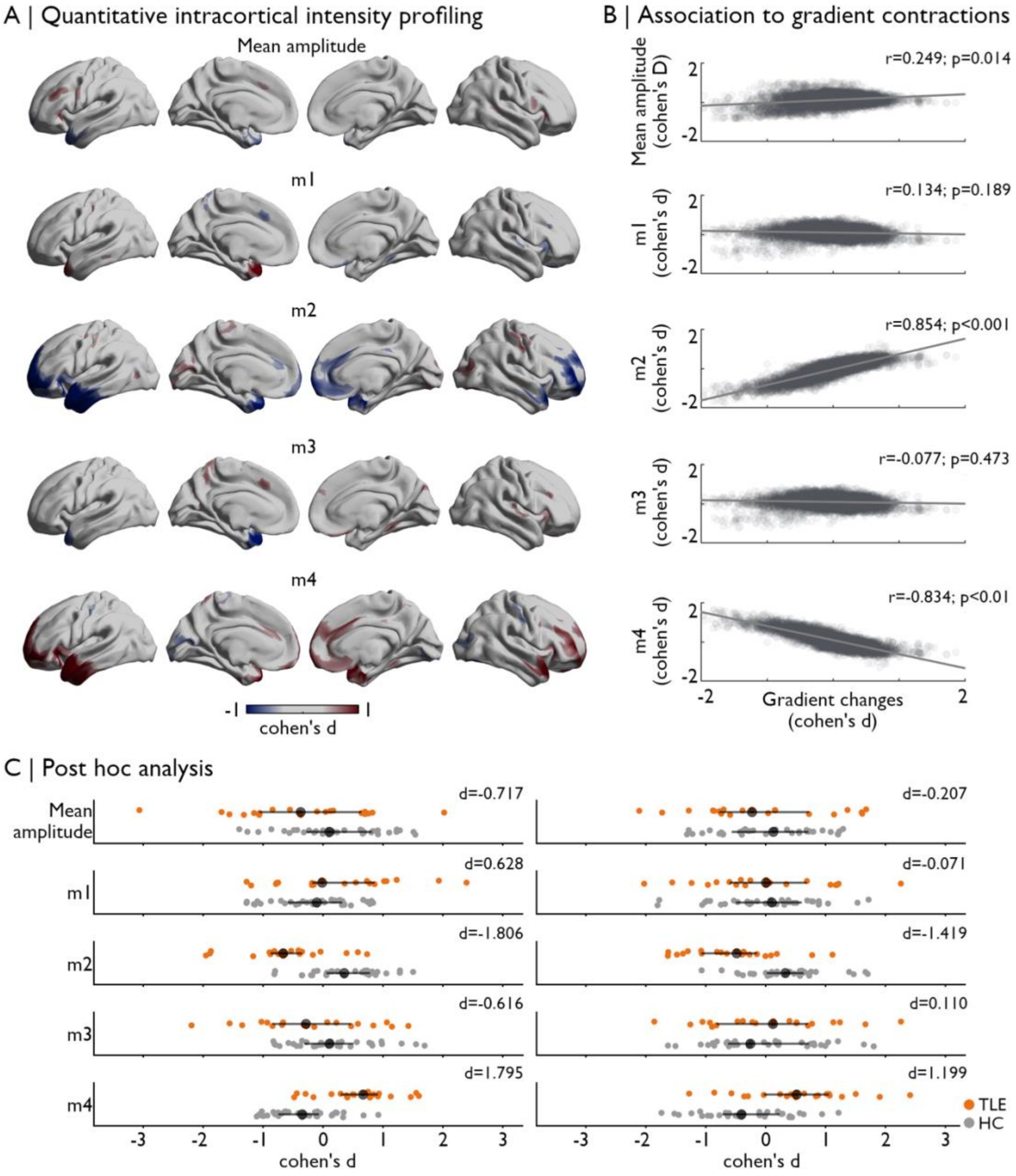


**Figure S5. Perturbed laminar microarchitecture underlies sensory-fugal gradient reorganizations in TLE. (A)** The shape of intracortical R1 profiles was quantified using the mean amplitude and four central moments, calculated across cortical depths at each vertex. After correction for age and sex, differences between TLE and HC groups could be seen across all moments (only effect sizes > 0.5 and < -0.5 are shown) **(B)** Spatial correlations between central moment and gradient change maps showed strongest cortex-wide correlations with m2 (standard deviation) and m4 (kurtosis). Statistical significance was determined using nonparametric spin permutation tests. **(C)** *Post hoc* analyses targeting significant clusters of gradient reductions highlighted more severe perturbations in laminar architecture in ipsilateral temporopolar relative to prefrontal regions. Larger black points and associated lines on each subplot represent median as well as 25th and 75th percentile of group distributions.

**FIGURE S6**

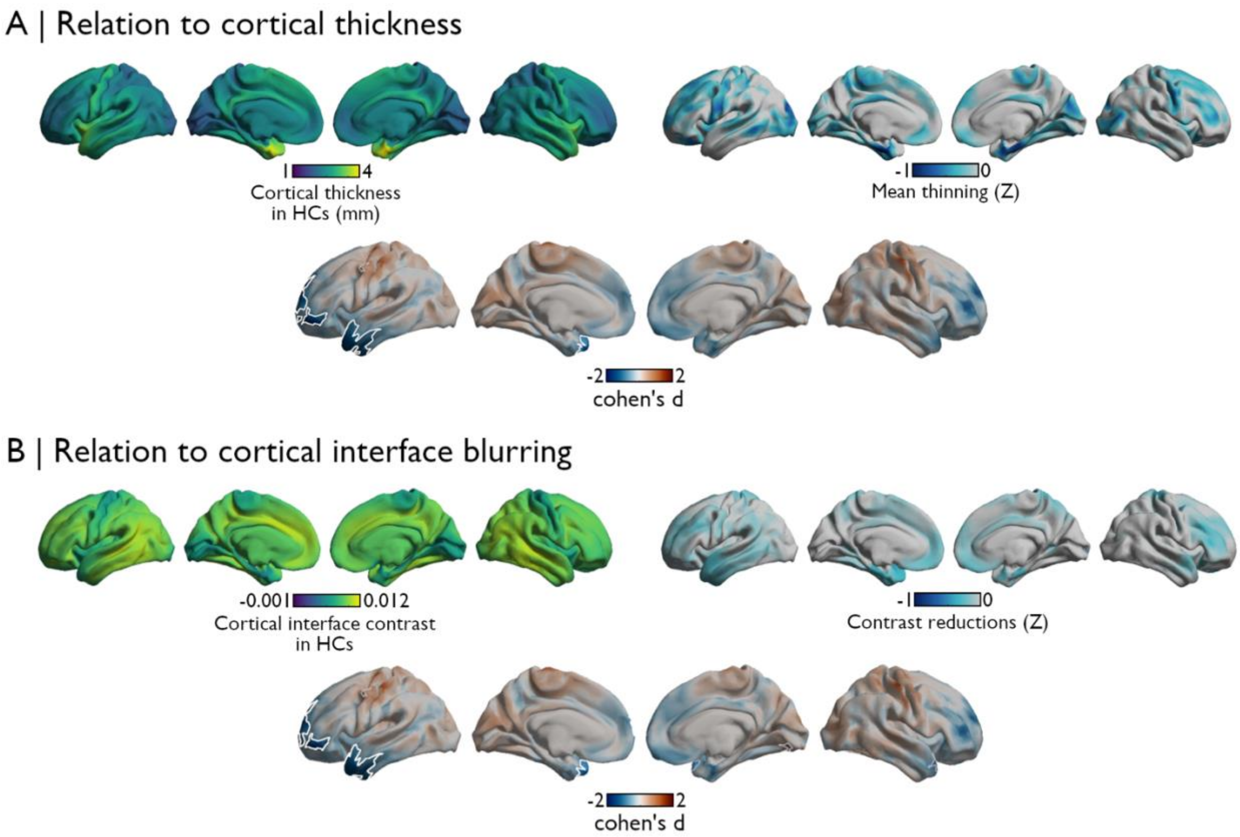


**Figure S6. Microstructural gradient reorganizations in TLE are robust to altered cortical thickness and blurring. (A)** Patients showed widespread and bilateral patterns of cortical thinning relative to controls, with stronger effects seen in mesio-temporal, frontal, and occipital cortices. Group comparisons of microstructural gradient scores controlling for vertex-wise cortical thickness were consistent with initial findings. **(B)** Subtle reductions in gray/white matter interface contrast were seen bilaterally in temporal and frontal regions in TLE. This metric captured distance-dependent differences in R1 intensity between superficial white matter and deep intracortical intensities. Group comparisons of microstructural gradient scores controlling for vertex-wise contrast alterations were also consistent with initial findings.

**FIGURE S7**

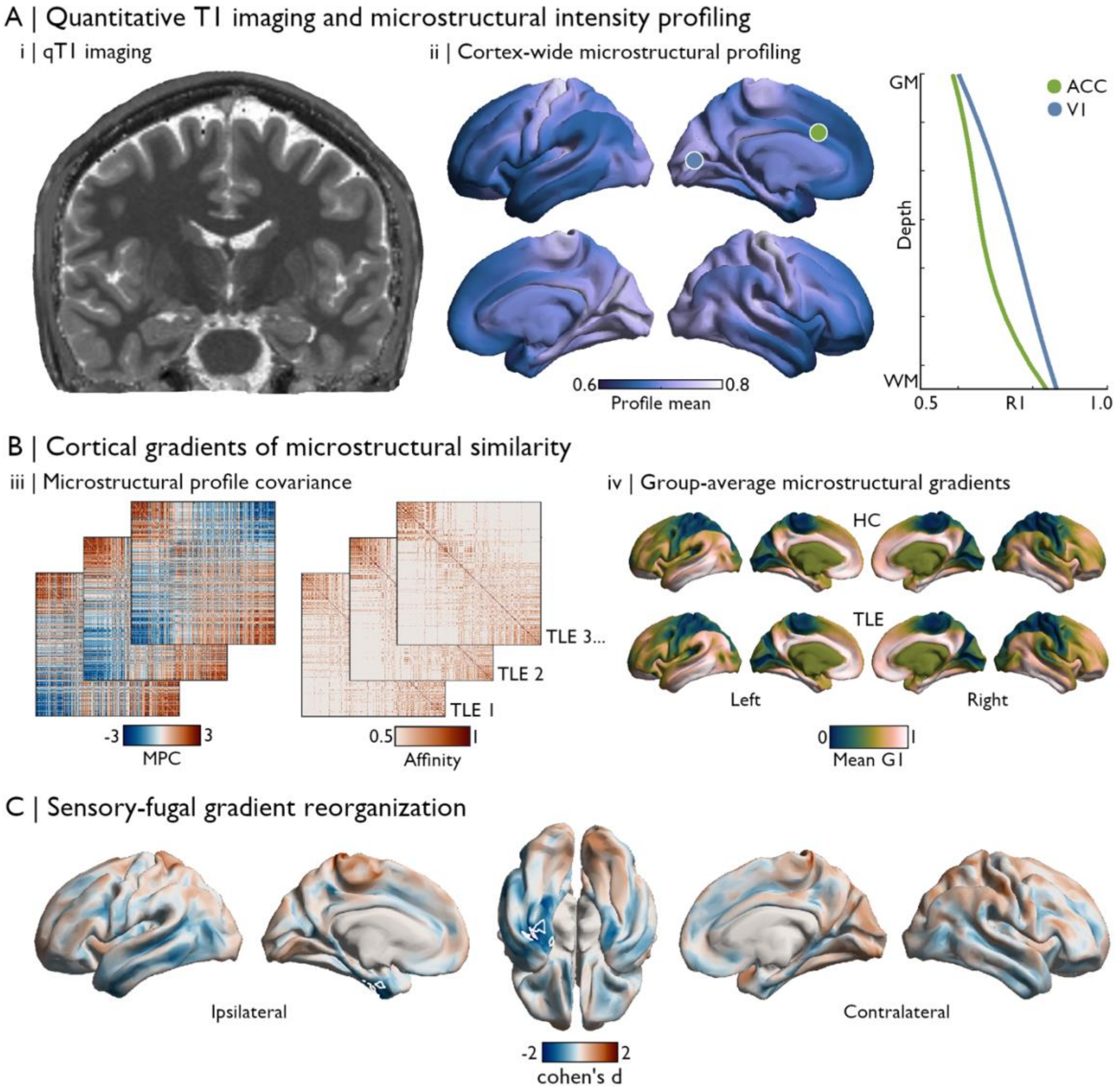


**Figure S7. Replicability of microstructural gradient changes in validation cohort. (A)** As in the discovery cohort, quantitative T1 imaging was used as a proxy of intracortical myeloarchitecture in TLE and HC individuals *(i)*. Intensities were sampled at 14 cortical depths between the pial and white matter boundaries, yielding vertex-wise microstructural intensity profiles. Mean R1 (1/T1) intensity calculated across cortical depths is displayed on the surface template. Profiles sampled in the primary visual cortex (V1, blue) and anterior cingulate cortex (ACC, green) showed different shapes in the validation cohort *(ii)*. **(B)** MPC matrices were constructed by cross-correlating vertex-wise intensity profiles using partial correlations controlling for the average cortex-wide profile, and normalized angle affinity matrices were generated from corresponding subject-level MPC matrices (*iii*). We applied diffusion map embedding to identify eigenvectors (gradients) describing main axes of variance in inter-regional similarity of cortical microstructural patterns. G1 is shown in (*iv*) for HC and TLE groups. Subject-specific gradients were aligned to the previously computed template derived from the discovery cohort data. **(C)** Surface-based linear models controlling for age and sex revealed significant differences in gradient scores between TLE and HC groups. Patients showed significant reductions in gradient scores in ipsilateral mesial and inferior anterior temporal lobe regions ($p_{FWE}$<0.001).

**FIGURE S8**

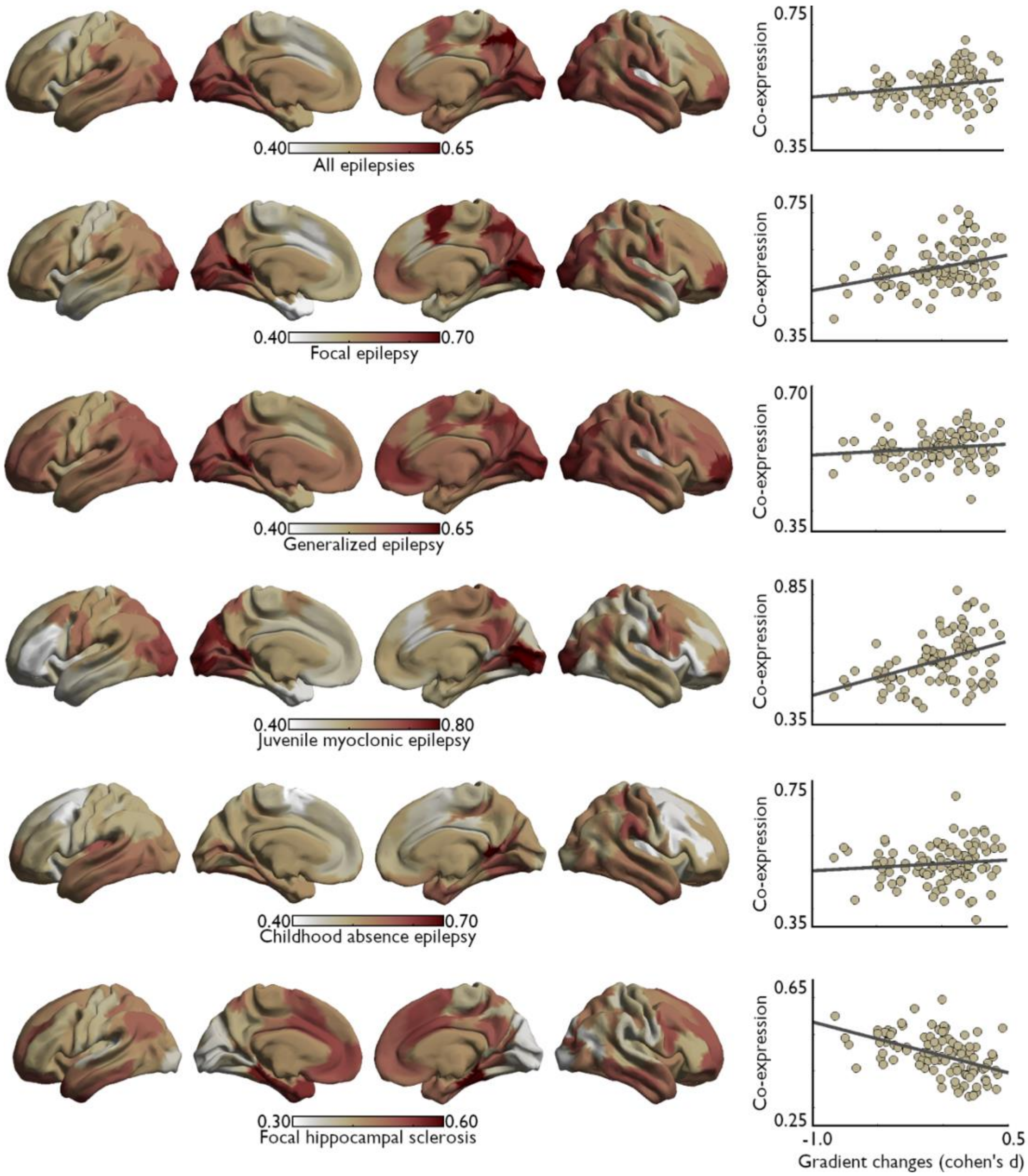


**Figure S8. Epilepsy syndrome gene co-expression maps and relationship to TLE-related microstructural gradient reorganizations.** Cross-referencing results of recent genome-wide association (GWAS) studies with data from the Allen Human Brain Atlas (AHBA) provided distinct gene co-expression patterns for several epilepsy syndromes (left). Only the correlation between co-expression patterns of genes associated with focal hippocampal sclerosis and microstructural gradient reorganizations seen in our TLE sample reached statistical significance after controlling for shared spatial autocorrelation. Scatter plots (right) depict the spatial relationship between each epilepsy syndrome (y-axis) and gradient changes (x-axis).

**FIGURE S9**

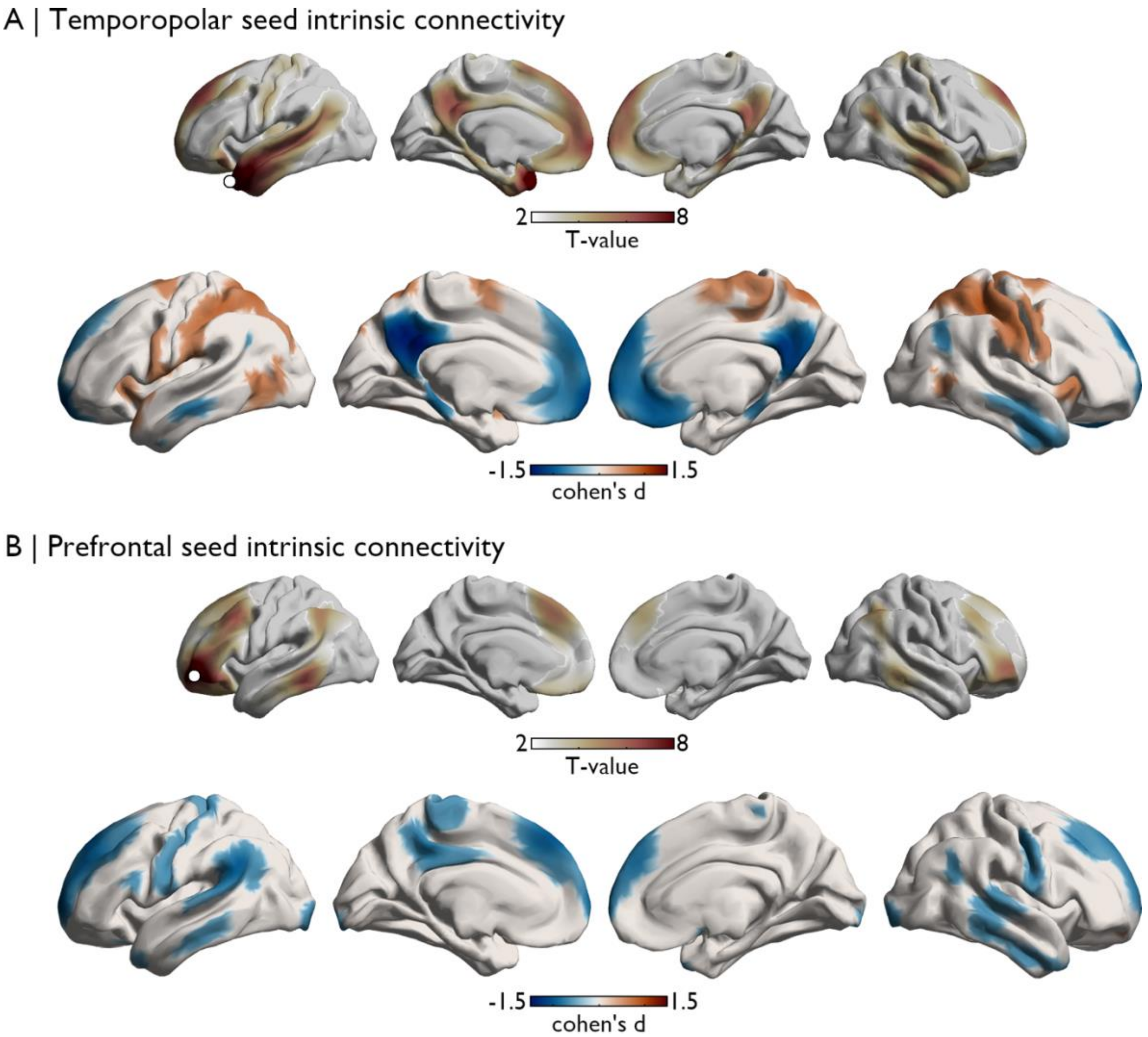


**Figure S9. Intrinsic functional connectivity changes associated with microstructural gradient seeds**. **(A)** Results of seed-based functional connectivity analysis during a resting-state acquisition, centered on peak regions in the temporopolar cluster of gradient contractions. Connectivity patterns in HCs are displayed in the top portion of the panel. Compared to HCs, TLE patients showed reduced connectivity to nodes of the default mode network, with strongest effects in the ipsilateral precuneus, as well as distributed connectivity increases with the anterior insula, lateral occipital regions, and fronto-parietal regions. (B) Results of seed-based functional connectivity analysis during a resting-state acquisition, centered on peak regions in the prefrontal cluster of gradient contractions. Connectivity patterns in HCs are displayed in the top portion of the panel. Compared to HCs, TLE patients showed reduced connectivity to distributed fronto-parietal regions, as well as lateral temporal cortices. In both A and B, Cohen's D maps are thresholded to show effect sizes larger than 0.5 and lower than -0.5. Connectivity between both seeds did not significantly differ between patients and controls in ipsilateral ($t$=-1.266; $d$=-0.434; $p$=0.212) and contralateral ($t$=-0.963; $d$=-0.334; $p$=0.340) hemispheres.

### **REFERENCES**

1. Garbelli R, Milesi G, Medici V, *et al*. Blurring in patients with temporal lobe epilepsy: clinical, high-field imaging and ultrastructural study. *Brain*. 2012;135(8):2337-2349.
2. Bernhardt BC, Fadaie F, de Wael RV, *et al*. Preferential susceptibility of limbic cortices to microstructural damage in temporal lobe epilepsy: A quantitative T1 mapping study. *Neuroimage*. 2018;182:294-303.
3. Hong S-J, Bernhardt BC, Schrader D, Caldairou B, Bernasconi N, Bernasconi A. MRI-based lesion profiling of epileptogenic cortical malformations. Springer; 2015:501-509.
4. Consortium TILAE. Genome-wide mega-analysis identifies 16 loci and highlights diverse biological mechanisms in the common epilepsies. *Nature communications*. 2018;9
5. Hawrylycz MJ, Lein ES, Guillozet-Bongaarts AL, *et al*. An anatomically comprehensive atlas of the adult human brain transcriptome. *Nature*. 2012;489(7416):391-399.
6. Larivière S, Paquola C, Park B-y, *et al*. The ENIGMA Toolbox: multiscale neural contextualization of multisite neuroimaging datasets. *Nature Methods*. 2021;18(7):698-700.
